# Comparative toxicogenomics of glyphosate and Roundup herbicides by mammalian stem cell-based genotoxicity assays and molecular profiling in Sprague-Dawley rats

**DOI:** 10.1101/2021.04.12.439463

**Authors:** Robin Mesnage, Mariam Ibragim, Daniele Mandrioli, Laura Falcioni, Eva Tibaldi, Fiorella Belpoggi, Inger Brandsma, Emma Bourne, Emanuel Savage, Charles A Mein, Michael N Antoniou

## Abstract

Whether glyphosate-based herbicides (GBHs) are more potent than glyphosate alone at activating cellular mechanisms, which drive carcinogenesis remains controversial. As GBHs are more cytotoxic that glyphosate, we reasoned they may also be more capable of activating carcinogenic pathways. We tested this hypothesis by comparing the effects of glyphosate with Roundup GBHs both *in vitro* and *in vivo*. First, glyphosate was compared with representative GBHs namely MON 52276 (EU), MON 76473 (UK) and MON 76207 (USA) using the mammalian stem cell-based ToxTracker system. Here, MON 52276 and MON 76473, but not glyphosate and MON 76207, activated oxidative stress and unfolded protein responses. Second, molecular profiling of liver was performed in female Sprague-Dawley rats exposed to glyphosate or MON 52276 (both at 0.5, 50, 175 mg/kg bw/day glyphosate) for 90 days. MON 52276 but not glyphosate increased hepatic steatosis and necrosis. MON 52276 and glyphosate altered the expression of genes in liver reflecting TP53 activation by DNA damage and circadian rhythm regulation. Genes most affected in liver were similarly altered in kidneys. Small RNA profiling in liver showed decreased amounts of miR-22 and miR-17 from MON 52276 ingestion. Glyphosate decreased mir-30 while miR-10 levels were increased. DNA methylation profiling of liver revealed 5,727 and 4,496 differentially methylated CpG sites between the control and glyphosate and MON 52276 exposed animals respectively. Apurinic/apyrimidinic DNA damage formation in liver was increased with glyphosate exposure. Altogether, our results show that Roundup formulations cause more biological changes linked with carcinogenesis than glyphosate.

## Introduction

Glyphosate is the most widely used herbicide worldwide. Although the use of glyphosate remained limited during the first two decades after its market introduction in 1974, worldwide use increased exponentially from the 1990s due to the rapid adoption of agricultural practices involving cultivation of transgenic soybeans, maize, cotton, sugar beet and other crops genetically engineered to tolerate applications of glyphosate formulated products such as Roundup (Benbrook 2016). In addition, glyphosate-based herbicides (GBHs) have increasingly been sprayed shortly before harvest to cause crop desiccation allowing a more rapid harvest, and for weed control in amenity and industrial areas.

The toxicity of glyphosate on human health and the environment is a controversial topic (Mesnage and Zaller 2021). The intensity in the debate on glyphosate toxicity increased dramatically when the World Health Organization’s International Agency for Research on Cancer (IARC) classified this compound as a probable (Class 2A) human carcinogen (Guyton et al. 2015). Although the existence of a direct link between glyphosate exposure and carcinogenesis in human populations at typical low exposure levels still remains to be established, IARC identified evidence in the peer reviewed scientific literature that glyphosate and commercial GBHs cause oxidative stress and DNA damage in animal and *in vitro* model systems (Guyton et al. 2015). However, findings from genotoxicity studies were often contradictory. Among the assays reviewed by IARC to evaluate possible genotoxic mechanisms, 77% reported positive results (Benbrook 2019).

An important knowledge gap, which complicates the debate on glyphosate carcinogenicity, is whether co-formulants included in GBHs could amplify glyphosate adverse effects. It was estimated that more than 750 different commercial herbicide formulations contained glyphosate in the US in 2015 (Guyton et al. 2015). These GBHs contain ingredients other than glyphosate, which also have toxic properties in humans (Sawada et al. 2011). The exposure to MON 0818 (a mixture of co-formulants) was identified as a worker safety issue during the 1980s because it was linked to occupational eye and skin injuries (Blondell 1986). Generally, although co-formulants are also toxic, long-term testing is not a regulatory requirement prior to their commercial authorization, and their composition remains company proprietary, confidential information. Thus, co-formulant toxicology and human exposure health effects remain a major gap in the regulation of these compounds (Mesnage and Antoniou 2018). The main surfactant present in GBHs, ethoxylated tallowamines, also known as POEA, was banned in the EU in 2016. However, it is not known if the more recent generations of GBHs containing co-formulant surfactants that are claimed to be less toxic, can still cause adverse health effects at doses which glyphosate alone is considered safe.

The mechanism or mechanisms by which GBHs may be carcinogenic are not fully known. Recent studies have provided mechanistic details suggesting a link between mitochondrial dysfunction caused by glyphosate and an increase in production of reactive oxygen species leading to DNA damage. Glyphosate has been known to affect the function of mitochondria by interference with the respiratory chain since the end of the 1970s (Olorunsogo et al. 1979, Olorunsogo and Bababunmi 1980, Olorunsogo 1990). More recently, in a study of *Caenorhabditis elegans* (Bailey et al. 2018), it was found that the glyphosate formulation TouchDown® Hitech inhibited mitochondrial respiration, resulting in a decreased proton gradient integrity and an inhibition of ATP production. Mitochondrial dysfunction induced by glyphosate was further confirmed by a study in zebrafish, *Danio rerio*, in which behavioural impairments induced by a GBH were linked to the effects of glyphosate on mitochondrial complex enzymatic activities and its resulting increased production of reactive oxygen species (Pereira et al. 2018). Studies on human peripheral blood mononuclear cells also suggest that glyphosate and its formulated products could generate DNA strand-breaks, and purine and pyrimidine oxidation via oxidative stress (Wozniak et al. 2018). Our previous study also suggested that glyphosate effects on the gut microbiota could be a source of oxidative stress (Mesnage et al. 2021), which implies that DNA damage from glyphosate exposure could arise from mechanisms which would not be detectable in commonly used in vitro systems.

However, the large number of glyphosate genotoxicity studies with contradictory results and divergent interpretations suggests that some uncontrolled factors are influencing the identification of genotoxic properties (Benbrook 2019). Since carcinogen hazard identification and risk assessment is increasingly relying on mechanistic data (Smith et al. 2016), we hypothesized that the use of the most recent generation of genotoxicity experimental systems can better inform as to whether exposure to glyphosate will activate mechanisms known to be key characteristics of carcinogens. As GBHs are more cytotoxic that glyphosate, we reasoned they may also be more capable of activating carcinogenic pathways. In order to test this hypothesis and the need to compare the toxicity of glyphosate with its commercial formulations, and to evaluate whether these products can active mechanisms of carcinogenicity, or cause DNA damage, we performed two sets of experiments.

First, we used a mammalian stem cell-based genotoxicity experimental system (ToxTracker Assay), which is designed to discriminate a chemical’s ability to induce DNA damage, oxidative stress and protein unfolding (Hendriks et al. 2016). This ToxTracker assay system was used to provide insight into the mechanisms underlying the toxicity of glyphosate and 3 commercial GBHs, namely the representative EU commercial herbicide formulation Roundup MON 52276, a US formulation MON 76207 (Roundup PROMAX) and a UK formulation MON 76473 (Roundup ProBio).

Second, we performed a subchronic study for 90-days in female rats administered with glyphosate (0.5, 50, 175 mg/kg bw/day), or MON 52276 at the same glyphosate equivalent doses. MON 52276 was chosen because it is the representative EU formulation and thus a suitable model for glyphosate herbicide hazard assessment. This duration was chosen in order to reflect the recommendations of OECD Guideline 408 (Repeated Dose 90-Day Oral Toxicity Study in Rodents), one of the most commonly used pesticide toxicity bioassays, which we complemented with a molecular phenotyping approach to potentially improve its sensitivity and predictivity of negative health outcomes. We combined high-throughput molecular profiling to highlight any alterations in the epigenome (DNA methylation) and transcriptome profile in liver (Figure 1). We also profiled small RNAs (∼20–30 nucleotides), which have emerged as a promising biomarker to track the development of disease (Harrill et al. 2016). We focused on the liver as it is a major target of injury by oxidative stress and inflammation (Cichoz-Lach and Michalak 2014). An increasing number of studies have revealed non-target effects of glyphosate on liver function (Chan and Mahler 1992, Benedetti et al. 2004, Beuret et al. 2005, Mesnage et al. 2017, Milić et al. 2018, Pandey et al. 2019, Mills et al. 2020). Some of these studies describe molecular phenotypes overlapping those found in cases of fatty liver disease, and which were associated with oxidative damage (Mesnage et al. 2017, Pandey et al. 2019, Mills et al. 2020). The acceptable daily intake of glyphosate in the European Union (EU) between 2002 and 2017 was based on liver damage detected after long-term administration of 60 mg/kg bw/day of glyphosate in rats (Commission 2002). These findings attest to the suitability of liver as an appropriate target organ to evaluate glyphosate and GBH carcinogenicity.

**Figure 1.**
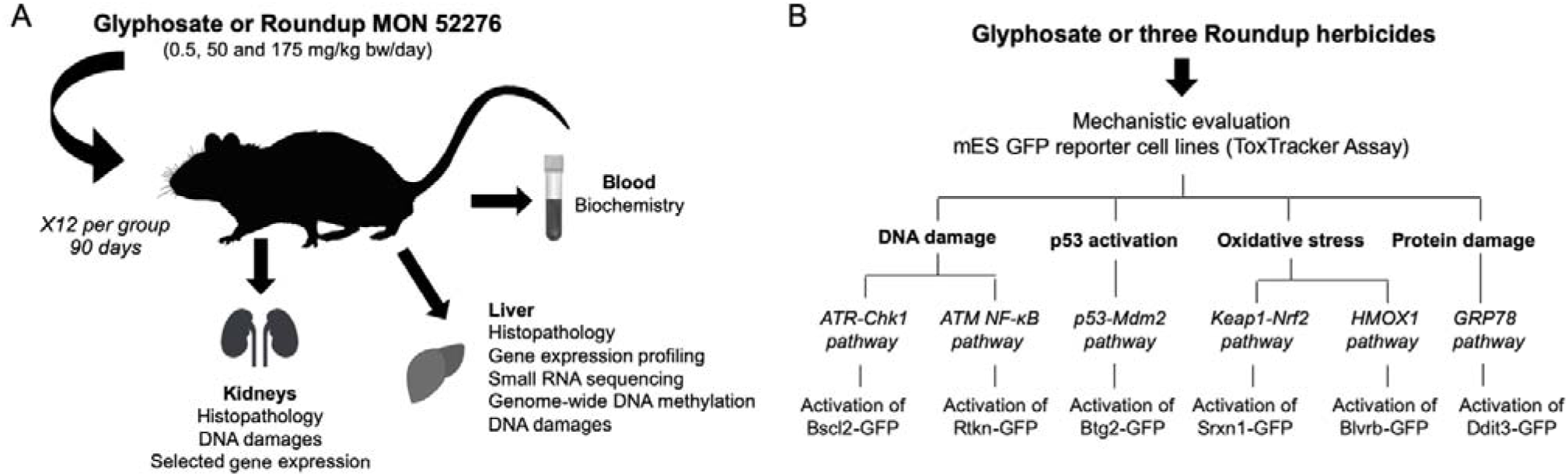
Experimental design. **A.** Female Sprague-Dawley rats were exposed to three concentrations of glyphosate or MON 52276 at the same glyphosate equivalent doses over 90-days. Molecular profiling techniques were applied in conjunction with standard histopathology and blood biochemistry analyses. **B.** The ToxTracker assay system was used to identify the potential carcinogenic mechanisms of glyphosate and three formulated glyphosate products (MON 52276, MON 76473 and MON 76207).

## Material and methods

### Experimental animals

The experiment was conducted with young adult female Sprague-Dawley rats at the Cesare Maltoni Cancer Research Center, Ramazzini Institute, (Bentivoglio, Italy) in accordance with Italian law regulating the use and humane treatment of animals for scientific purposes (Decreto legislativo N. 26, 2014. Attuazione della direttiva n. 2010/63/UE in materia di protezione degli animali utilizzati a fini scientifici. - G.U. Serie Generale, n. 61 del 14 Marzo 2014). The experiment was authorised by the ad hoc commission of the Italian Ministry of Health (authorization N. 447/2018-PR).

Female Sprague-Dawley rats were generated in-house following an outbreeding programme at the Cesare Maltoni Cancer Research Center, Ramazzini Institute, (Bentivoglio, Italy). Animals were tagged using ear punch marking using the Jackson Laboratory system. Female animals were chosen in order to follow-up our previous studies, where we provided evidence suggesting that the long-term exposure to Roundup GT Plus caused the development of fatty liver disease (Mesnage et al. 2015, 2017) and gut microbiome and blood metabolome disturbances (Mesnage et al. 2021). Animals were randomised after weaning to have at most one sister per litter in each group and to have a homogeneous body weight across groups. Rats were housed in polycarbonate cages (41x25x18 cm) with stainless steel wire tops and a shallow layer of white wood shavings as bedding. The animals were housed 3 per cage. They were kept at a temperature of 22±3°C and relative humidity of 50±20%, with a 12-hour light/dark cycle. Cages were periodically rotated to minimise effects due to cage position.

Treatment of animals was as previously described (Mesnage et al. 2021). In brief, groups of 12 female Sprague-Dawley rats of 8 weeks of age were administered for 90 days with glyphosate and MON 52276 as Roundup BioFlow (Italy) at the same glyphosate equivalent dose via drinking water of 0.5 mg, 50 mg and 175 mg/kg body weight per day (mg/kg bw/day), which respectively represent the EU acceptable daily intake (ADI), the EU no-observed adverse effect level (NOAEL) and the US NOAEL (European Food Safety 2015). A group of 12 animals not subjected to the glyphosate treatments was used as control. The EU NOAEL of 50 mg/kg bw/day is the lowest NOAEL dose among animal studies performed to determine glyphosate toxicity and derive an ADI of 0.5 mg/kg bw/day. In the US, the equivalent NOAEL is 175 mg/kg bw/day, which was used to derive a chronic Reference Dose (cRfD) of 1.75 mg/kg bw/day. The daily water and food consumption per cage were measured before the start of the experiment, and weekly for the entire 13-week duration of the treatment. The body weight of animals was measured weekly during the study period. In the group treated with the highest dose of Roundup, the concentration of MON 52276 had to be reduced starting from 6^th^ week of treatment because of the lower solution consumption of the animals. The mean daily intake of glyphosate in animals of this group, for the whole period of treatment, was therefore 130 mg/kg bw/day instead of the 175 mg/kg bw/day.

Animals were checked for general status three times a day, seven days a week, except for non-working days when they were checked twice. No deaths occurred during the study period.

### Histopathology

Prior to sacrifice, animals were anesthetised by inhalation of a 70% CO_2_/30% O_2_ mixture. All sacrificed animals were subjected to complete necropsy. Liver and kidneys were alcohol-fixed, trimmed, processed and embedded in paraffin wax. Sections of 3-6 μm were cut for each specimen of liver and kidneys and stained with haematoxylin and eosin. All slides were evaluated by a pathologist and all lesions of interest were re-evaluated by a second pathologist. The histopathological nomenclature of lesions adopted were classified according to the international nomenclature INHAND (International Harmonization of Nomenclature and Diagnostic Criteria) and RITA (Registry of Industrial Toxicology Animal Data).

### Biochemistry

At the time of sacrifice, approximately 7.5 ml of blood was collected from the *vena cava* which were centrifuged at 3000rpm for 10 minutes to obtaining serum. Serum biochemistry analysis was performed at IDEXX BioAnalytics (Stuttgart, Germany), a laboratory accredited to ISO 17025 standards. Sodium and potassium levels were measured by indirect potentiometry. Albumin was measured by a photometric Bromocresol green test. Alkaline phosphatase ALP was measured by IFCC with the AMP-buffer method, glucose by Enzymatic UV-Test (Hexokinase method), cholesterol by Enzymatic colour test (CHOD-PAP), blood urea nitrogen by enzymatic UV-Test, gamma-glutamyl-transferase by Kinetic colour test International Federation of Clinical Chemistry (IFCC), aspartate and alanine aminotransferase by kinetic UV-test (IFCC+ pyridoxal-5-phosphate), creatinine by kinetic colour test (Jaffe’s method), lactate dehydrogenase (LDH) by the IFCC method, and triglycerides using an enzymatic colour test (GPO-PAP) on a Beckman Coulter AU 480 instrument.

### DNA and RNA extraction

DNA and RNA were extracted from tissues taken at the time of sacrifice and which had been stored at -80°C. DNA and RNA extraction, cDNA library preparation and sequencing for transcriptome profiling was performed as described previously (Mesnage et al. 2021). Liver and kidneys from animals exposed to a dose of 50 mg/kg bw/day of glyphosate or to MON 52276 at the same glyphosate equivalent dose were used. In brief, genomic samples extracted from rat liver displayed a high molecular weight with DINs (DNA integrity score) ranging from 7.5 to 10, and average concentrations from 370 ng/µL. All liver RNA samples had integrity numbers (RIN) ≥ 7 as verified with the Agilent 2100 Bioanalyser (Agilent Technologies, Waldbronn, Germany). A260/A280 were ≥ 1.8 as verified with Thermo Scientific™ NanoDrop 2000 spectrophotometer (ThermoFisher Scientific, UK). RNA was extracted from ∼30 mg rat liver or kidney (from the cortical area) using the All Prep DNA/RNA/miRNA Universal Kit (Qiagen, Hilden, Germany) according to manufacturer’s instructions. All kidney RNA samples had A260/A280 ratios ≥ 1.7 as verified with Thermo Scientific™ NanoDrop 2000 spectrophotometer (ThermoFisher Scientific, UK).

### Transcriptomics

The preparation of cDNA libraries was performed using 100 ng total RNA using the NEBNext® Poly(A) mRNA Magnetic Isolation Module, the NEBNext® Ultra™ II Directional RNA Library Prep Kit, and indexed with NEBNext® Multiplex Oligos for Illumina® (96 Index Primers) (New England Biolabs, Ipswich, Massachusetts, USA). Sample libraries at 1.1 pM were sequenced twice on the NextSeq500, and 75 bp paired-end reads were generated for each library using the Illumina NextSeq®500 v2.5 High-output 150 cycle kit (75bp paired-end reads, Illumina Inc., Cambridge, UK). A total of 455,308,427 reads (average of 12,647,456 ± 2,461,518 reads per sample) were generated for the 36 liver samples.

### Reduced representation bisulfite sequencing

A total of 100 ng of total genomic liver DNA was diluted and processed using the Premium Reduced Representation Bisulfite Sequencing (RRBS) Kit (Diagenode, Denville, NJ, USA) as per the manufacturer’s instructions. Libraries were sequenced to 75 base pair single end on a NextSeq 500 (Illumina, CA, USA). Data was aligned to the rat reference genome Rn6 with Bismark (Krueger and Andrews 2011). A total of 650,358,053 reads (average of 18,065,501 ± 4,846,025 reads per sample) were generated.

### DNA damage assay

DNA damage was measured in the liver and kidneys of rats exposed to a dose of 50 mg/kg bw/day of glyphosate or to MON 52276 at the same glyphosate equivalent dose. We used a quantitative method to measure the formation of apurinic/apyrimidinic (AP) sites, which are common DNA lesions caused by oxidative damage. This was done using the DNA Damage Assay Kit (AP sites, Colorimetric) (Abcam, ab211154) (Abcam plc, Cambridge, UK) according to manufacturer’s instructions.

### RT-qPCR

We selected 5 genes indicative of oxidative damage to DNA from the transcriptomics of liver and measured their expression in kidney samples from rats exposed to a dose of 50 mg/kg bw/day of glyphosate or to MON 52276 at the same glyphosate equivalent dose. Extracted RNA was retrotranscribed to cDNA using the SuperScript™ VILO™ cDNA Synthesis Kit (ThermoFisher Scientific, Loughborough, UK). A total of 100 ng cDNA was then amplified using TaqMan™ assays for *Nr1d2* (Rn00596011_m1), *Gadd45g* (Rn01352550_g1), *Nr1d1* (Rn01460662_m1), *Ier3* (Rn03993554_g1), *Clec2g* (Rn01462968_m1), using *Actb* (Rn00667869_m1) as an internal reference standard. Reactions were conducted as technical duplicates, with a TaqMan™ Fast Advanced Master Mix (ThermoFisher Scientific, UK) on the Applied Biosystems QuantStudio 6 Flex Real-Time quantitative PCR System. The delta-delta Ct method (Livak and Schmittgen 2001) was used to calculate the relative gene expression of the target genes in kidney samples.

### Small RNA profiling

We created miRNA libraries from 100 ng total RNA using the QIAseq miRNA library kit (Qiagen, Hilden, Germany) according to manufacturer’s instructions. This kit was chosen to incorporate unique molecular indices (UMIs) into each cDNA to enable correction for PCR bias. Library concentrations were measured using Qubit™ dsDNA HS Assay Kit (Thermo Fisher, USA). Size of the libraries and calculation of the final molarity for sequencing was performed with the High Sensitivity D1000 ScreenTape Assay on an Agilent 2200 TapeStation System (Agilent Technologies, Waldbronn, Germany). Average size of the fragments was 181 ± 22 bp, with TapeStation molarity of 31.8 ± 12.9 nM. Libraries were normalised to 3nM. All samples and all libraries were of good quality and 75 bp single end reads were generated for each library using the Illumina NextSeq®500 v2.5 High-output 150 cycle kit (Illumina Inc., Cambridge, UK). A total of 153,887,020 reads were generated for the 36 samples (4,274,639 ± 1,905,806 reads per sample).

### ToxTracker® Assay

The ToxTracker® assay system is a panel of six validated green fluorescent protein (GFP) gene-based mouse embryonic stem (mES) reporter cell lines that can be used to identify the biological reactivity and potential carcinogenic properties of newly developed compounds in a single test (Hendriks et al. 2016). The ToxTracker mES cell lines contain a GFP reporter gene inserted into genes whose expression is known to increase in response to various carcinogenic stimuli (oxidative stress, DNA damage, protein unfolding). The biomarker gene *Bscl2* informs on DNA damage following activation of ATR/Chk1 DNA damage signaling, while *Rtkn* is a marker of NF-kB signaling reflecting double-strand DNA breaks. Oxidative stress is evaluated with 2 genes, with *Srxn1* reflecting an Nrf2 antioxidant response and *Blvrb* reflecting an Nrf2 independent response. Protein damage is evaluated as the unfolded protein response by measuring *Ddit3.* Activation of the p53 tumor suppressor response pathway is evaluated with *Btg2.* This assay was performed as previously described (Hendriks et al. 2016). We compared the genotoxic properties of glyphosate and MON 52276, and also two other formulated GBH products: MON 76207 (Roundup PROMAX on the US market) and MON 76473 (Roundup ProBio on the UK market).

In brief, glyphosate, Roundup BioFlow (MON 52276), MON 76207 and MON 76473 were diluted to a glyphosate equivalent concentration of 50 mM in water and the pH adjusted to 7.2. First, wild type mES cells (strain B4418) were exposed to 20 different concentrations of the test substances to determine cytotoxic effects and select 5 concentrations to be tested in the six independent ToxTracker mES reporter cell lines. Induction of GFP reporter gene expression was determined after 24 h exposure using flow cytometry using a Guava EasyCyte Flow cytometer (Luminex). Potential metabolic activation was included in the ToxTracker assay by addition of S9 liver extract from aroclor1254-induced rats (Moltox). Cells were exposed to five concentrations of the test samples in the absence and presence of 0.25% S9 extract and required co-factors (RegenSysA+B, Moltox) for 24 h. Positive reference treatments with cisplatin (DNA damage), diethyl maleate (oxidative stress), tunicamycin (unfolded protein response) and aflatoxin B1 (metabolic activation of progenotoxins by S9) were included in all experiments. Solvent concentration was the same in all wells and never exceeded 1% DMSO.

### Statistical analysis

ToxTracker assay data analysis was performed using ToxPlot, a dedicated data analysis software package created to import raw green fluorescent protein (GFP) reporter gene expression data from the flow cytometry measurements, calculate GFP induction levels and cytotoxicity, and perform statistical analysis of the data. The ToxTracker assay is considered to give a positive response when a compound induces at least a 2-fold increase in GFP expression in any of the six cell assay systems. Only GFP induction at concentrations that do not cause more than 75% cytotoxicity are used for the ToxTracker analysis. A total of three independent repeat experiments in either the absence or presence of metabolic activation of the test substances and the control compounds cisplatin, diethyl maleate, tunicamycin and aflatoxin B1 were performed.

All the 84 animals (7 groups of 12 animals) were subjected to an analysis of food and water consumption, body and organ weights, a serum biochemistry analysis, and a histopathological evaluation of the kidneys and liver. We focused our mRNA-Seq, small RNA-Seq, reduced representation bisulfite sequencing, and DNA damage analysis on 36 samples from the cohort of rats exposed to the EU NOAEL (50 mg/kg bw/day) for MON 52276 and glyphosate, as well as the untreated controls.

Food and water consumption, as well as body and organ weights, were analysed with Kruskal-Wallis test with Dunnett’s post hoc comparisons with STATA 10 software. Incidence of non-neoplastic lesions was evaluated with a Fisher’s exact test (one and two-tailed; one-sided results were also considered, since it is well established that only an increase in incidence can be expected from the exposure, and incidence in the control group are almost always 0). The analysis of linear trend, for incidence of pathological lesions, was obtained using the Cochran-Armitage trend test (OECD 2011, Shockley and Kissling 2018). Statistical significance for the serum biochemistry data was evaluated using a One-way ANOVA with post-hoc Tukey HSD (honestly significant difference).

The mRNA-seq data was analysed with Salmon (Patro et al. 2017). This tool was used to quantify transcript abundance by mapping the reads against a reference transcriptome (Ensembl Release Rattus Norvegicus 6.0 cDNA fasta). The Salmon output was then imported into R version 4.0 (Team 2019) using the Bioconductor package tximport. We created a transcript database containing transcript counts, which was used to perform a differential gene expression analysis using DESeq2 (Love et al. 2014). Genes with less than 5 counts in 5 samples were removed. Unnormalized raw counts were used to create the DESeq2 model. Tables of outcome data were generated using the DESeq2 results function to create a results table with log2 fold changes, Wald test p value and adjusted p values for multiple comparisons made according to the Benjamini-Hochberg procedure. The contrast argument of results function was then used to make pair-wise comparisons between untreated and treated samples. Differentially expressed genes were identified by DESeq2 with a cut-off of more than 1.5-fold-change and an adjusted p-value of less than 0.05. We finally used goseq to perform a gene ontology of biological processes and a Kyoto encyclopedia of genes and genomes (KEGG) pathway enrichment analysis accounting for transcript length biases using the gene annotations included in the R package org.Rn.eg.db 3.11.4 (Young et al. 2010). The relationship between the different significant GO terms was visualised using the REVIGO web server (Supek et al. 2011)

DNA methylation calls from RRBS data were extracted with Bismark (Krueger and Andrews 2011). The output from Bismark was then imported into R and analysed with Methylkit (Akalin et al. 2012). DNA methylation calls were annotated using RefSeq gene predictions for rats (rn6 release) with the package genomation (Akalin et al. 2014). Other annotations were retrieved using the genome wide annotation for rat tool org.Rn.eg.dbR package version 3.8.2. Statistical analysis was performed with logistic regression models fitted per CpG using Methylkit functions. P-values were adjusted to Q-values using the SLIM method (Wang et al. 2011).

Small RNA sequencing data was performed with miRge3.0 (Patil and Halushka 2021). The MiRge3.0 tool was run with Pyton 3.8 in a Conda environment with bowtie v1.3.0, cutadapt v2.6, Samtools v1.9 and RNAfold v2.4.14. The miRge library was the rat specific small-RNA annotation library provided as part of the miRge3.0 tool. Low-quality bases (Q<20) as well as uncharacterised bases were trimmed from the 5’ and/or 3’ ends of each read. We also used miRge3.0 to predict new miRNAs. The minimum and maximum length of the retained reads for novel miRNA detection was 16 and 25 bp, respectively. Minimum read counts to support novel miRNA detection was 2. We took into account UMIs in order to remove PCR duplicates and used standard small RNA libraries as a reference (Patil and Halushka 2021). The output of miRge3.0 was then imported in R (v4.0.0) for statistical analysis. The miRNA counts were used to perform a differential gene expression analysis using DESeq2 in a similar manner as described above for the mRNA-seq data (Love et al. 2014). TargetScan 7.1 was used to predict biological targets of miRNAs among glyphosate-altered mRNAs by searching for the presence of conserved sites that match the miRNA seed region (Lewis et al. 2005). The resulting miRNA-mRNA network was visualised using the R package for network visualization visNetwork.

We ultimately analysed the different datasets of liver molecular profiles with an orthogonal partial least squares discriminant analysis (OPLS-DA) to evaluate the predictive ability of each omics approach. OPLS-DA can be used to distinguish the variability corresponding to the experimental perturbation from the portion of the data that is orthogonal; that is, independent from the experimental perturbation. The R package ropls version 1.20.0 was used with a nonlinear iterative partial least squares algorithm (NIPALS) (Thévenot et al. 2015). Prior to analysis, experimental variables were centred and unit-variance scaled. Since PLS-DA methods are prone to overfitting, we assessed the significance of our classification using permutation tests (permuted 1,000 times).

### Data availability

All the raw data is made available in Dryad (https://doi.org/10.5061/dryad.1g1jwstxk). The study metadata describing the different identifiers linking the different analyses (Supplementary File 1), as well as data for water consumption (Supplementary File 2), food consumption (Supplementary File 3), body weights (Supplementary File 4), pathology (Supplementary File 5) and serum biochemistry (Supplementary File 6) is made available. The raw data for the liver transcriptomics is available at the GEO accession number GSE157426. Transcriptome count data (Supplementary File 7), and gene expression statistics (Supplementary File 8 and 9), GO enrichments (Supplementary File 10 and 11) and KEGG pathway enrichments (Supplementary File 12 and 13), are available for MON 52276 and glyphosate, respectively. The raw data for the small RNA profiling is available at the GEO accession number GSE182559. Quantification files from Salmon are available (Directory transcriptomes). Small RNA count data (Supplementary File 14), predicted new miRNAs (Supplementary File 15), as well as summary statistics for MON 52276 (Supplementary File 16) and glyphosate groups (Supplementary File 17) are also provided. Count data for all unique molecular indices are provided (Directory smallRNA). Ct values for the expression of genes in kidneys is provided as Supplementary File 18. The raw data from the RRBS analysis is available at the GEO accession number GSE157551. Methylation summary statistics are available for MON 52276 (Supplementary File 19) and glyphosate (Supplementary File 20). Bismark files with the number of C bases that are methylated or unmethylated at each genomic coordinate are provided (Directory methylation). We also provide the results of the DNA damage assay for each animal (Supplementary File 21). The code used to perform the statistical analysis is made available as supplementary data (Supplementary File 22).

## Results

The primary aim of this investigation was to compare toxicogenomic properties of glyphosate and representative members of its commercial Roundup formulations. The ToxTracker assay system was first used to test if glyphosate and 3 GBHs activate mechanisms, which are known to be key characteristics of carcinogenesis. We also performed an *in vivo* animal experiment comparing standard histopathology and serum biochemistry measures to high-throughput molecular profiling of epigenome (DNA methylation) and transcriptome (mRNA and small RNA) profiles in liver.

### Oxidative stress and unfolded protein responses

The correct functioning and accuracy of the ToxTracker assay was demonstrated by exposure to various reference substances and monitoring expression of the GFP reporter genes, which are known to increase in expression in response to various carcinogenic stimuli (Figure S3). The genotoxic compound cisplatin showed induction of a DNA damage response (*Bscl2* and *Rtkn* induction) and p53-mediated cellular stress (*Btg2* induction). Diethyl maleate induced primarily the oxidative stress related reporters *Srxn1* and *Blvrb*, whilst tunicamycin induced an unfolded protein stress response (*Ddit3* induction). The positive control compound aflatoxin B1, which requires metabolic activation to become genotoxic, selectively induced the *Bscl2* and *Rtkn* reporters when tested in the presence of S9 metabolic liver extract (Figure S3).

Cytotoxicity of the different GBHs varied markedly (Figure S2). The formulation MON 76207 was highly cytotoxic (LC50 ∼ 107 µM) compared to MON 52276 (LC50 ∼ 2.7 mM) and MON 76473 (LC50 ∼ 1.8 mM). Glyphosate caused no cytotoxic effects at a concentration of 5 mM in the absence or presence of the metabolising S9 liver extract (Figure S2). Figure 2 summarises the results of the ToxTracker assay for glyphosate and the three different GBHs (MON 52276, MON 76473, MON 76207). Induction in expression of the GFP reporter genes is shown in the absence (Figure 2A to 2D) or presence of S9 liver extracts (Figure 2I to 2L). We also report cell survival in absence (Figure 2E to 2H) or presence of S9 liver extract (Figure 2M to 2P). The two reporters reflective of oxidative stress are Srxn1-GFP and Blvrb-GFP. Induction of the Srxn1-GFP reporter is associated with activation of the Nrf2 antioxidant response and activation of the Blvrb-GFP reporter is associated with the Hmox1 antioxidant response. Activation of the oxidative stress reporter Srxn1-GFP was observed with MON 76473 (Figure 2B, 2J) and MON 52276 (Figure 2D, 2L). MON 76207 weakly activated (>1.5-fold) the Srxn1-GFP reporter only in the presence of S9 metabolic liver extract. The Ddit3-GFP reporter, associated with protein damage and an unfolded protein response, was activated upon exposure to MON 76473 (Figure 2B, 23J) and MON 52276 (Figure 2D, 2L) in the absence and presence of S9 liver extract. No activation of a DNA damage response (via *Bscl2* and *Rtkn* induction) and p53-mediated cellular stress (via *Btg2* induction) was observed with any of the test compounds.

**Figure 2.**
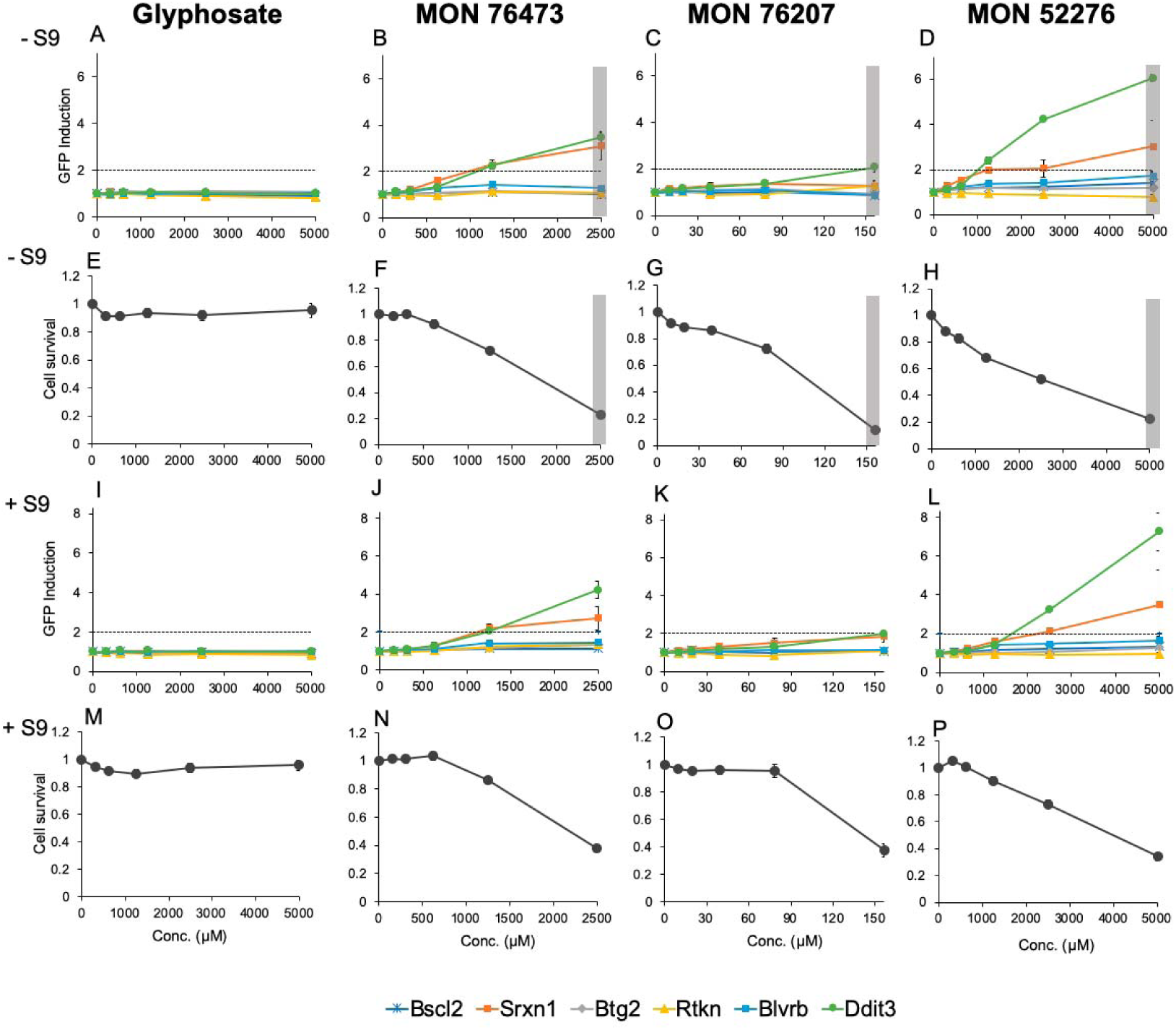
GBHs activated oxidative stress and unfolded protein responses in the ToxTracker GFP-based mouse embryonic stem (mES) reporter cell line system. The six ToxTracker mES reporter cell lines were used to detect oxidative stress (Srxn1 and Blvrb), protein damage and an unfolded protein response (Ddit3), the activation of a DNA damage response (Bscl2 and Rtkn) and p53-mediated cellular stress (Btg2). **Panels A to D:** induction of reporter gene expression in absence of S9 liver extracts and associated changes in cell survival **(panels E to H)**. **Panels I to L:** induction of the reporter gene expression in the presence of S9 liver extract and cell survival **(panels M to P)**. Grey shaded area denotes a concentration at which cell survival is less than 25% and thus no reliable interpretation regarding reporter gene induction from the treatment can be ascertained from this dose. n = 3 independent repeat experiments.

Collectively, these ToxTracker assay results show that Roundup herbicides activated oxidative stress and unfolded protein responses while glyphosate did not.

### Roundup MON 52276 caused more serious liver lesions than glyphosate

Rats were exposed to glyphosate and its representative EU Roundup formulation MON 52276. The doses used were chosen as they represent the EU ADI (0.5 mg/kg bw/day), the EU NOAEL (50 mg/kg bw/day) and the US NOAEL (175 mg/kg bw/day). No significant differences were observed in either feed consumption (Table S1) nor in mean body weight (Table S2), and only the group administered with the highest dose of MON 52276 (175 mg/kg bw/day glyphosate equivalent dose) had a reduced water consumption (Table S3).

All treated and control animals were subjected to a histopathological evaluation of the kidneys and liver (Figure 3). No lesions were detected in liver of control animals whereas a low frequency (2/8 animals) of pelvic mineralisation, inflammation and epithelial pelvic necrosis was observed in their kidneys in the control animals (Table 1). There was a statistically significant increase of animals bearing all types of lesions (p=0.036; Fisher two-tail test) and in particular hyaline casts in kidneys (p=0.0037; Fisher two-tail test) in the group exposed to 0.5 mg/kg bw/day glyphosate. Inflammation was the most frequent lesion observed in the kidneys, with non-statistically significant increased incidence in all treated groups compared to controls, except for the group treated with 50 mg/kg of glyphosate (Table 1, Glyphosate columns). Contrastingly, a dose-dependent and statistically significant increase in the incidence of liver lesions (fatty liver changes, necrosis) was observed in rats treated with MON 52276 (Table 1, MON 52276 columns; Figure 3). A statistically significant increase of animals bearing necrosis and all types of lesions in the group treated with 175mg/kg bw/day of MON 52276 (p=0.047; Fisher one-tail test) was detected. There was also a statistically significant linear trend in animals bearing fatty change (p=0.010; Cochran-Armitage test), necrosis [(p=0.002; Cochran-Armitage test), and all types of lesions (p=0.002; Cochran-Armitage test) in MON 52276 treated groups. An increase in liver lesions was also detected in animals treated with glyphosate at all doses, although this did not reach statistical significance (Table 1, Glyphosate columns). Liver and kidney weights were unchanged (Table S4). Blood clinical biochemistry only revealed increased creatinine levels in the groups treated with glyphosate but not with MON 52276 (Figure 4).

**Figure 3.**
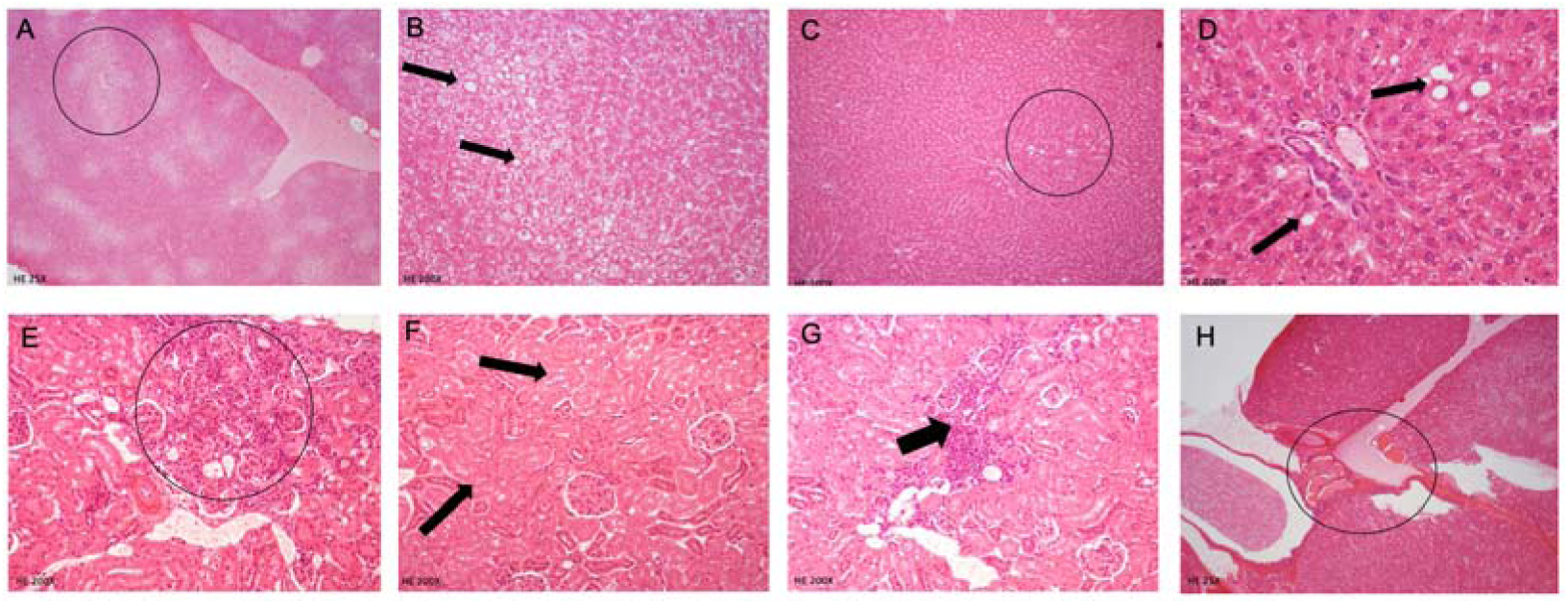
Pathological observations in rats following subchronic 90-day exposure to glyphosate and Roundup MON 52276. This selection of cases illustrates the identification and grading of histopathological observations. Diffuse and severe fatty change with necrosis of hepatocytes (circle and arrows) in animal N.132 treated with Roundup MON 52276 at a 175 mg/kg bw/day glyphosate equivalent dose **(A, B)**. Animal N.111 treated with MON 52276 at 50mg/kg bw/day with moderate focal areas of fatty change (circle and arrows) with necrosis of hepatocytes **(C, D)**. Kidney tubular fibrosis and inflammation (N. 26, glyphosate 0.5 mg/kg bw/day) **(E).** Kidney tubular degeneration (N. 47, glyphosate 50 mg/kg bw/day) **(F)**. Animal N. 32 treated with glyphosate (0.5 mg/kg bw/day) showing hyaline casts in tubules (circle) severe grade, in kidneys **(G)**.

**Figure 4.**
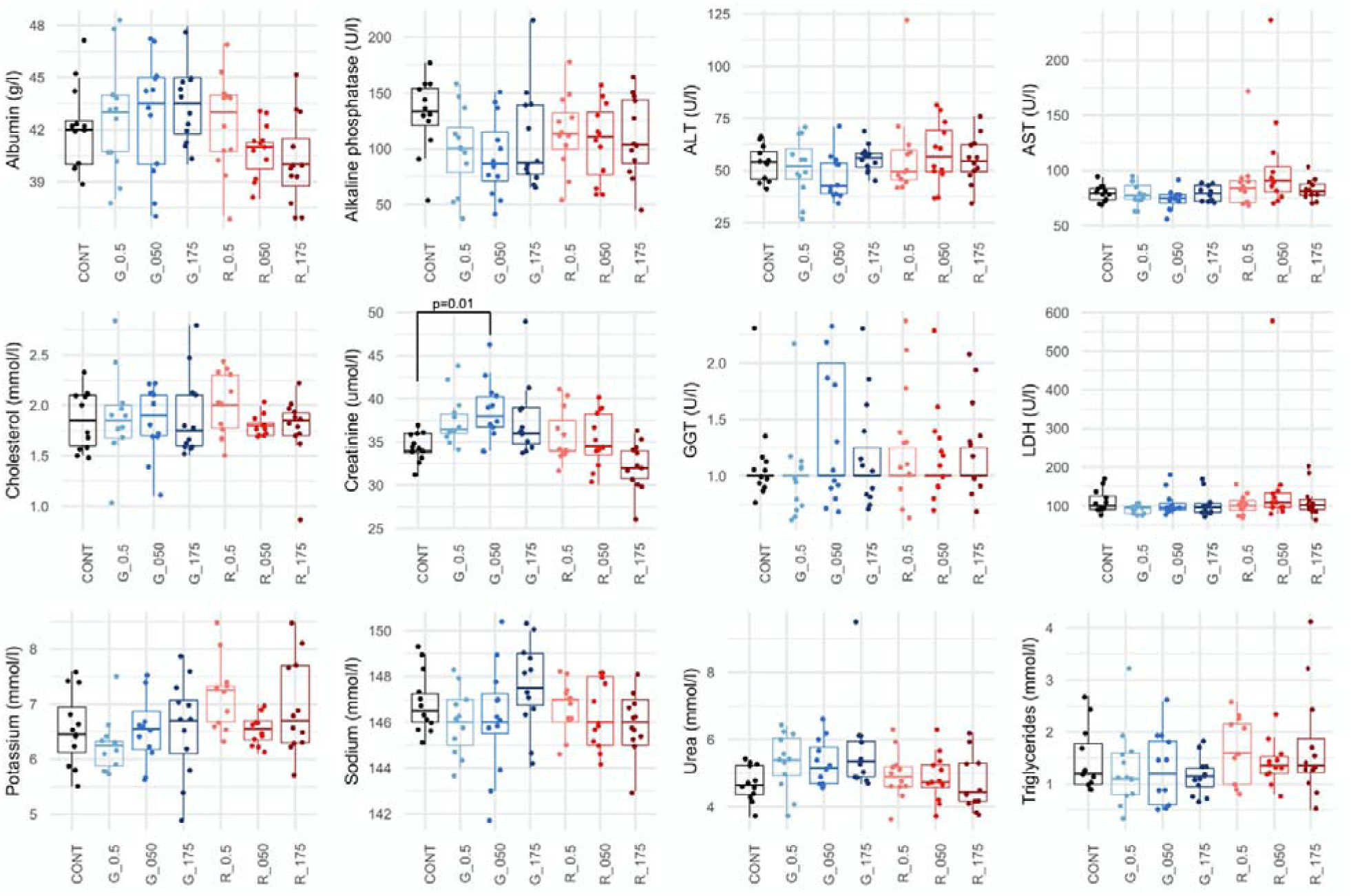
Serum biochemistry analysis of rats following subchronic 90-day exposure to glyphosate and Roundup MON 52276. Clinical biochemistry evaluation in female rats administered with glyphosate (G_0.5: 0.5 mg/kg bw/day; G_050: 050 mg/kg bw/day; G_175: 175 mg/kg bw/day;) and MON 52276 (R_0.5: 0.5 mg/kg bw/day; R_050: 050 mg/kg bw/day; R_175: 175 mg/kg bw/day). Results at the end of the treatment period revealed minor changes with only creatinine showing a statistically significant increase at the highest dose of glyphosate. Serum biochemistry values are shown as box plots with the median, two hinges (the 25th and 75th percentiles), and two whiskers extending to the furthest observation ≤ 1:5 times the interquartile range, along with individual values for each analyte (solid circles). n = 12 per group. The p-value is from a one-way ANOVA with post-hoc Tukey HSD.

**Table 1.**
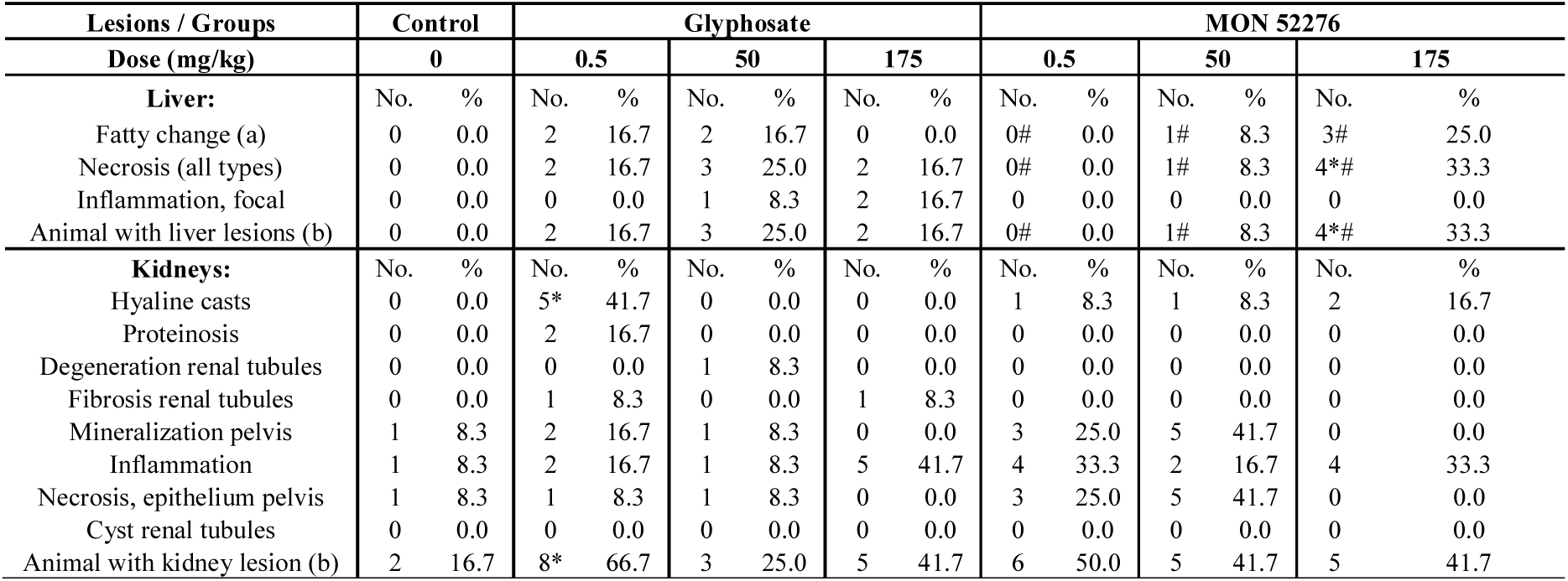
Histopathological evaluation of liver and kidneys in adult female Sprague-Dawley rats administered for 90-days with glyphosate and Roundup MON 52276. The table shows incidence of lesions in the liver and kidneys of all animals. (**a**) Fatty change includes from mild to severe lesions or associated to necrosis. (**b**) One animal could bear more than one lesion*. Statistically significance (P≤0.05) was assessed using the Fisher Exact test (one-tailed test); # p-values (P≤0.01) associated with the Cochran-Armitage test for trend. n = 12 per group.

### Liver transcriptomics reveals pathways of oxidative damage to DNA

In order to obtain insight into the possible effects of glyphosate and MON 52276 on liver function, we then used high-throughput molecular profiling techniques to search for molecular changes, which could act as predictors of the development of liver disease. We focused our analysis on liver samples from the cohort of rats exposed to the EU NOAEL (50 mg/kg bw/day). Animals exposed to 50 mg/kg bw/day were chosen because rats exposed to the highest dose of MON 52276 (175 mg/kg bw/day glyphosate equivalent dose) had a reduced water consumption indicating that a maximum palatable dose was reached. This could have confounded the results. In addition, the lowest level of exposure (0.5 mg/kg bw/day glyphosate) was not used in order to describe mechanisms at a dose at which there was a visible phenotypic effect (pathological signs observed with MON 52276).

Whilst a total of 20 genes had their expression altered (one gene was downregulated and 19 upregulated) by glyphosate (FDR = 5%) (Figure 5A), exposure to MON 52276 at the same glyphosate equivalent dose altered the expression of 98 genes (50 genes downregulated and 48 upregulated) (Figure 5B). This correlated well with the findings from the histopathology analysis (Table 1) showing that MON 52276 was causing more biological changes than glyphosate. Functional analysis failed to provide significant insights about the effects of glyphosate because of the low number of genes having their expression altered by this compound. However, important functional insights about the effects of MON 52276 were obtained. Using Gene Ontology (GO) annotations, we found that the exposure to MON 52276 affected the expression of genes associated with 173 biological processes (Figure 5D), with the most significant being the response to steroid hormones (adj.p = 0.0001) and radiation (adj.p = 0.002). A pathway analysis suggested that TP53 signaling (adj.p = 0.0008) and the regulation of circadian rhythms (adj.p = 0.01) were over-represented among the genes having their expression altered by MON 52276. Genes having their expression changed in relation to the TP53 signaling pathway were *Cdkn1a* (Cyclin Dependent Kinase Inhibitor 1A, FC = -2.5), *Gadd45a* (Growth Arrest And DNA Damage Inducible Alpha, FC = 1.7)*, Chek2* (Checkpoint Kinase 2, FC = 1.9), *Sesn2* (Sestrin 2, FC = -1.5) and *Gadd45g* (Growth Arrest And DNA Damage Inducible Gamma, FC = 2.5). The two most affected genes were *Nr1d1* (Nuclear Receptor Subfamily 1 Group D Member 1, FC = 8.8) and *Nr1d2* (Nuclear Receptor Subfamily 1 Group D Member 2, FC = 3.1). Fold changes in gene expression of the genes having their expression altered by MON 52276 were well correlated to those affected by glyphosate alone (Figure 5C, Pearson’s r = 0.83, p < 2.2e-16), suggesting that glyphosate affected the same pathways as MON 52276, even if only effects of MON 52276 reached statistical significance. Based on these results we were able to derive a short-list of genes whose expression was similarly altered by both glyphosate and MON 52276 and which could constitute a transcriptome signature for exposure to this class of herbicide (Table 2).

**Figure 5.**
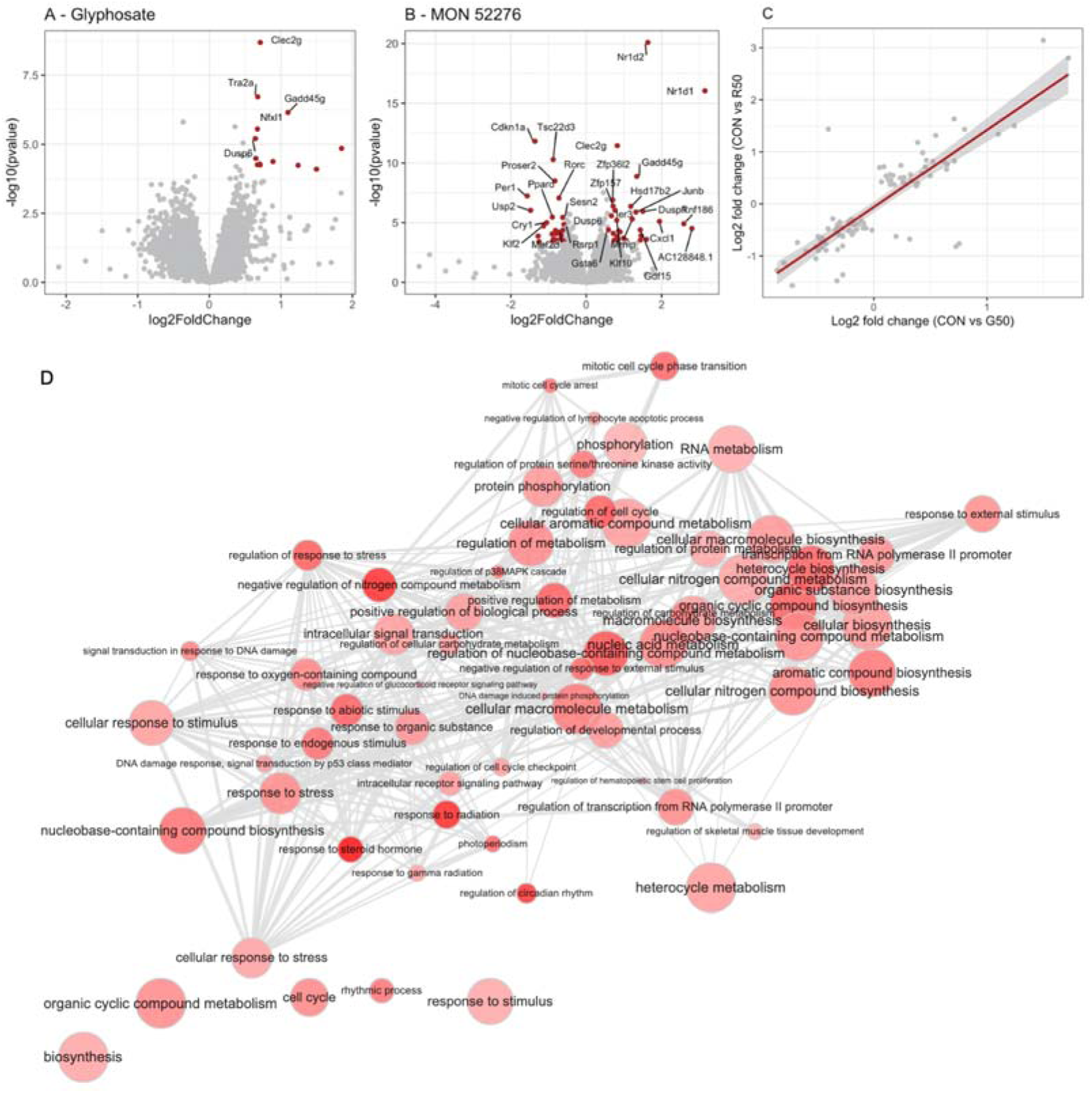
Transcriptome of liver from female Sprague-Dawley rats exposed for 90-days to glyphosate or Roundup MON 52276. A volcano plot showing the fold changes and statistical significance in the expression of genes affected by exposure to glyphosate **(A)** or MON 52276 **(B)** at a dose of 50 mg/kg bw/day. Genes were considered as differentially expressed (red dots) if their count was found to be statistically significant after an analysis with DeSeq2. Fold changes in gene expression caused by glyphosate and MON 52276 were well correlated **(C)**. GO terms enriched for the group of rats exposed to MON 52276 (nodes) and their relation (edges) are shown **(D)**. Bubble color indicates the significant adjusted p-value ranging from 0.05 to 0.001. n = 12 per group.

**Table 2.** Transcriptome signature in liver of female Sprague-Dawley rats exposed to both glyphosate and Roundup MON 52276. Genes whose expression was statistically significantly changed (adj-p < 0.05) after exposure to both glyphosate and Roundup MON 52276 were selected. FC, fold changes for glyphosate (gly) or MON 52276 (mon). n = 12 per group.

| Gene | Name | p gly | adjp gly | FC gly | p mon | adjp mon | FC mon |
| --- | --- | --- | --- | --- | --- | --- | --- |
| Clec2g | C-type lectin domain family 2 | 2.0x10<sup>-9</sup> | 2.0x10<sup>-5</sup> | 1.64 | 3.5x10<sup>-12</sup> | 1.0x10<sup>-8</sup> | 1.77 |
| Gadd45g | Growth Arrest and DNA Damage Inducible Gamma | 7.1x10<sup>-7</sup> | 2.3x10<sup>-3</sup> | 2.14 | 1.3x10<sup>-9</sup> | 2.6x10<sup>-6</sup> | 2.53 |
| C3 | Complement component 3 | 1.6x10<sup>-6</sup> | 3.8x10<sup>-3</sup> | -1.29 | 1.8x10<sup>-4</sup> | 2.9x10<sup>-2</sup> | -1.22 |
| Dusp6 | Dual specificity phosphatase 6 | 6.2x10<sup>-6</sup> | 8.6x10<sup>-3</sup> | 1.56 | 2.7x10<sup>-6</sup> | 1.4x10<sup>-3</sup> | 1.59 |
| Ier3 | Radiation-Inducible Immediate-Early Gene IEX-1 | 3.2x10<sup>-5</sup> | 2.6x10<sup>-2</sup> | 1.57 | 9.4x10<sup>-7</sup> | 6.9x10<sup>-4</sup> | 1.70 |
| Nr1d2 | Nuclear receptor subfamily 1 group D member 2 | 5.5x10<sup>-5</sup> | 3.3x10<sup>-2</sup> | 1.63 | 8.1x10<sup>-21</sup> | 9.5x10<sup>-17</sup> | 3.10 |
| Dusp1 | Dual-specificity phosphatase 1 | 5.7x10<sup>-5</sup> | 3.3x10<sup>-2</sup> | 2.36 | 1.1x10<sup>-6</sup> | 7.9x10<sup>-4</sup> | 2.83 |
| Zfp422 | Zinc finger protein 422 | 6.3x10<sup>-5</sup> | 3.4x10<sup>-2</sup> | 1.30 | 2.3x10<sup>-4</sup> | 3.5x10<sup>-2</sup> | 1.27 |
| Nr1d1 | Nuclear Receptor Subfamily 1 Group D Member 1 | 7.9x10<sup>-5</sup> | 4.0x10<sup>-2</sup> | 2.82 | 9.2x10<sup>-17</sup> | 5.4x10<sup>-13</sup> | 8.84 |

### Gene expression was altered by glyphosate or MON 52276 in kidneys

Transcriptome alterations caused by both glyphosate and MON 52276 in liver were marked by statistically significant changes in the expression of 9 genes (Table 2) with consistent fold changes (Figure 5A and 5B). We evaluated whether the expression of these genes could also be disturbed in kidneys to indicate DNA damage from glyphosate exposure (Figure 5D). In order to test this hypothesis, we measured expression of 5 genes from this list of 9 transcriptome biomarker candidates namely, *Nr1d1, Nr1d2*, *Clec2g, Ier3,* and *Gadd45g* in kidney, which showed the most significant alterations in liver samples (Table 2). Only *Nr1d2*, *Clec2g* and *Gadd45g* transcripts were detectable in kidney samples with differential expression between these genes found to be comparable to that observed in the liver samples (Figure 6). Overall, this suggested that kidneys and thus potentially other tissues may have been damaged by the exposure to MON 52276 and glyphosate.

**Figure 6.**
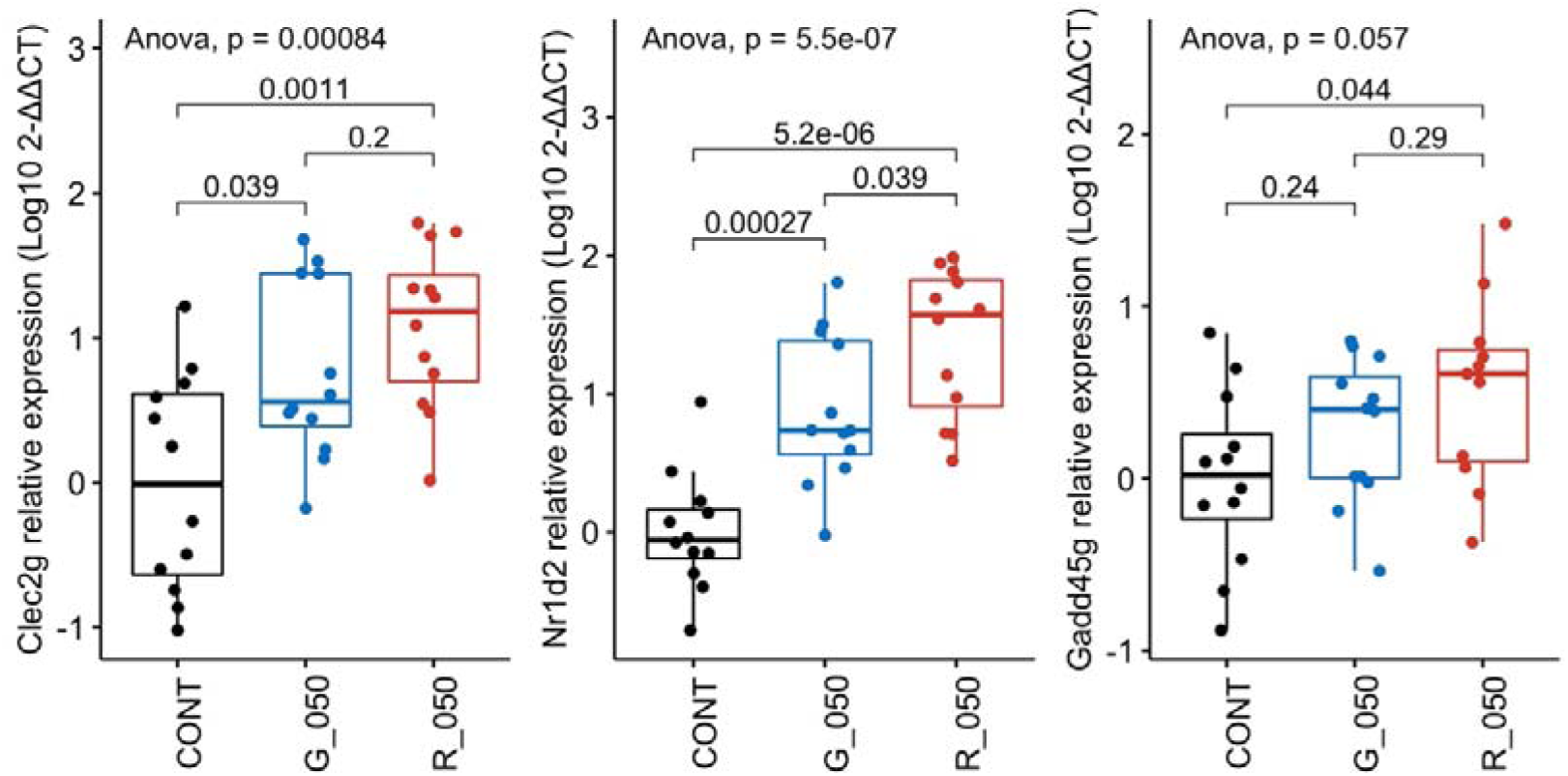
Expression of genes most significantly increased in liver by glyphosate and MON 52276 are similarly altered in kidneys. Total RNA was extracted from kidneys of rats exposed for 90 days to 50 mg/kg bw/day glyphosate (G_050) or Roundup MON 52276 (R_050) at the same glyphosate equivalent dose, and control animals ingesting regular tap water (CONT). Relative expression levels of *Nr1d2*, *Clec2g* and *Gadd45g* transcripts were determined by RT-qPCR and compared to and reported here as delta-delta Ct values, shown as box plots with the median, two hinges (the 25th and 75th percentiles), and two whiskers extending to the furthest observation ≤ 1:5 times the interquartile range, along with individual expression values for each gene (solid circles). n = 12 per group. The p-value is from a one-way ANOVA with post-hoc Tukey HSD.

### miRNA profiles suggests that regulation of gene expression by MON 52276 occurs partly post-transcriptionally

We also undertook a profiling of small RNAs in the same liver samples to assess if gene expression changes could be linked to this epigenetic system. An average of 49.7% of the total number of good quality reads were classified as miRNA while 30.4% were unclassified (Figure 7A). The remaining reads were tRNA (7.4%) and mRNA (9.8%). These included 677 unique miRNAs, which were identified in the miRNA database, and 9 novel predicted miRNAs. Statistical analysis showed that miR-22 and miR-17 had their levels decreased by exposure to MON 52276, whilst miR-30 had its levels decreased by glyphosate, and miR-10 had its levels increased by glyphosate. We then evaluated whether the mRNA, which have been found to have their levels altered were potentially regulated by these miRNAs (Figure 7B). The two miRs having their levels altered by MON 52276 (miR-22 and miR-17 levels) were predicted to regulate levels of 15 and 9 mRNAs respectively, which were also altered by MON 52276. None of the miRs affected by glyphosate treatment were predicted to regulate glyphosate-altered mRNAs. This suggests that the alterations in gene expression patterns by MON 52276 (Figure 5B, Table 2) occur at least in part at the post-transcriptional level mediated by changes in miRs.

**Figure 7.**
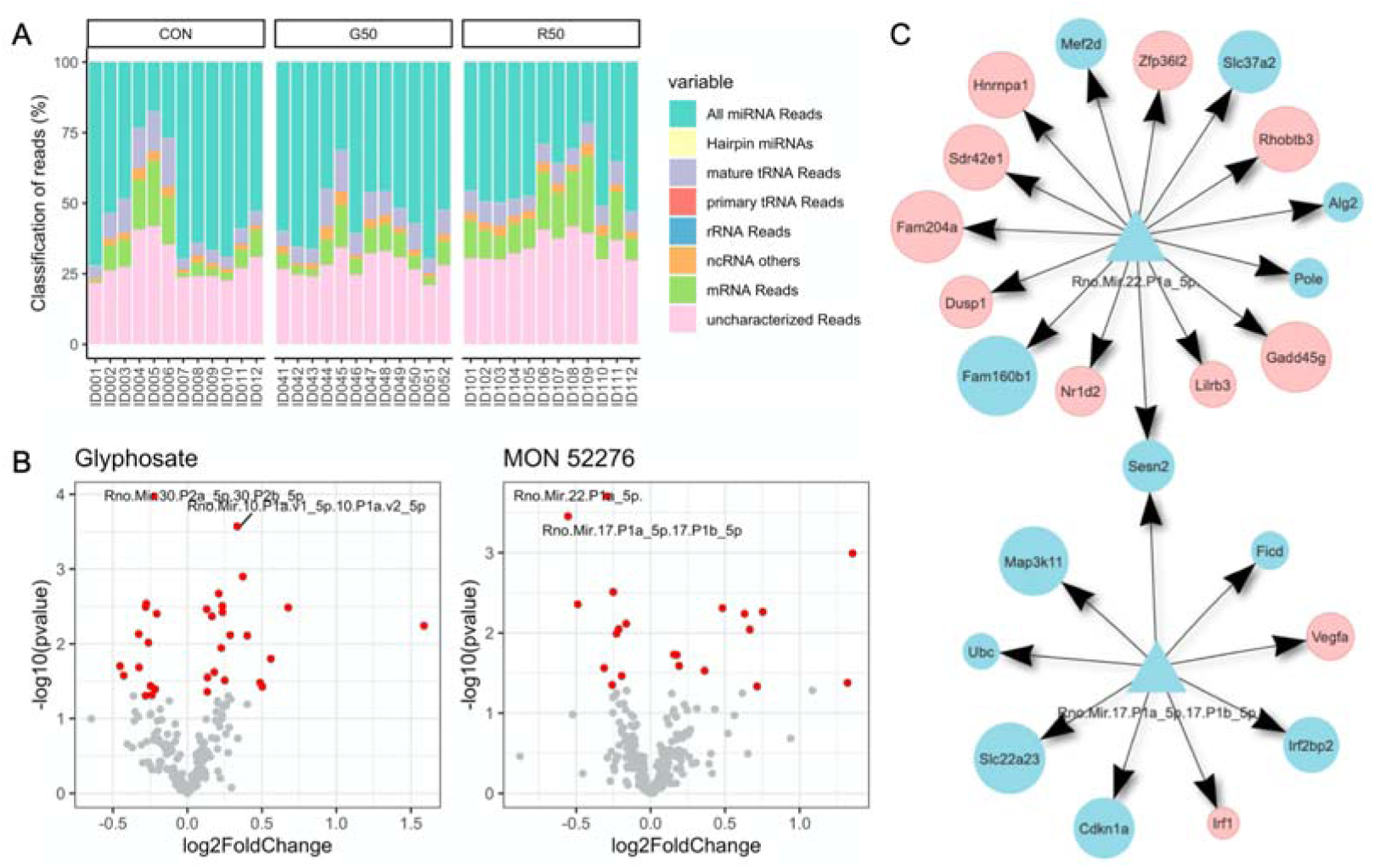
Small RNA profiles in liver samples from rats exposed to glyphosate or MON 52276. **A.** Small RNAs were classified based on their sequences using miRge3.0. **B.** Volcano plots showing the results from the analysis of miRNA differential abundance for glyphosate (left panel) or MON 52276 (right panel) exposed animals. Red dots represent miRNA with their unadjusted p-value < 0.05. Only 4 miRNAs were found to pass corrections for multiple comparisons (labelled dots). **C.** Genes predicted to be regulated by the differentially expressed miR-22 (top diagram) and miR-17 (lower diagram), which had their expression levels significantly decreased by MON 52276. TargetScan was used to predicts gene targets (circles) of miRNAs (triangles). Targets which were not found to have their expression levels affected by MON 52276 were filtered out. Genes are classified as downregulated (blue) or upregulated (red). None of the miRs affected by glyphosate treatment were predicted to regulate glyphosate-altered mRNAs. n = 12 per group.

### DNA methylation profiling reflected changes in the epigenome landscape

There was no difference in the percentage of methylated cytosines in CpG islands in either glyphosate or MON 52276 exposed rats (42.0 ± 1.6% for controls vs 42.6 ± 3.0% and 41.7 ± 1.7% for glyphosate and MON 52276 respectively). The distribution of the percentage DNA methylation per base was bimodal, as expected since a given C-residue is generally either methylated or not in a given cell (Figure 8A). CpG di-nucleotides near transcription start sites tend to be unmethylated (Figure 8B). Overall, we identified 5,727 and 4,496 differentially methylated CpG sites (DMS) (FDR < 0.05) with a modest methylation difference (> 10%) between the control group and the group of rats exposed to glyphosate and MON 52276, respectively (Figure 8C). When both treatments were compared, we found that 1100 CpG sites were found to be differentially methylated to the same degree in both glyphosate and MON 52276 administered groups. The amplitude of these changes ranged from -35 to 34%. Percentage methylation changes were very well correlated between glyphosate and MON 52276 treatment groups (Pearson’s r = 0.95, p < 2.2e-16), which suggests that they are not due to random variation (Figure 7D). Among these 1100 loci, DMS were in intergenic regions (50.7%) and introns (35.5%), and less within exons (8.4%) and at gene promoters (5.4%). DMS were located at an average distance of 59.8kb from transcriptional start sites (TSS).

**Figure 8.**
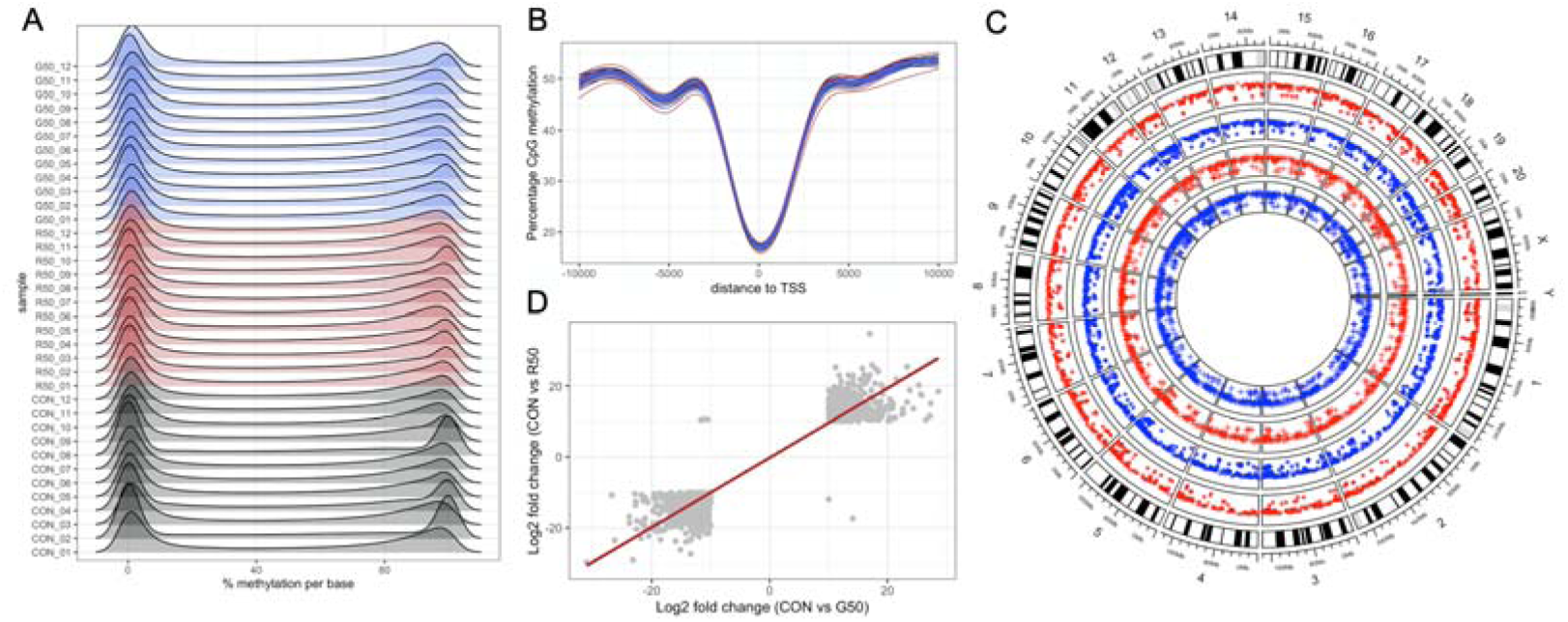
Reduced Representation Bisulfite Sequencing (RRBS) of the DNA from liver samples from female Sprague-Dawley rats exposed to glyphosate or its formulated product Roundup MON 52276. **A.** Distribution of percentage methylation per base **B.** CpG methylation around transcriptional start sites (TSS). **C.** Circos plot shows the location of differentially methylated CpG sites (blue track, hypomethylated; red track, hypermethylated; round symbols in outer tracks, glyphosate; cross symbols in inner tracks, MON 52276). **D**. Correlation of methylation changes for MON 52276 and glyphosate. n = 12 per group.

We evaluated if some DMS were present in the genomic regions surrounding the differentially expressed genes (DEGs) identified by RNA sequencing (Figure 5). No DMS were found in the genomic regions within 50 kb around the TSS of these genes. In addition, no gene had its expression changed among the differentially methylated CpG sites, which were attributed to promoters (Table 7). There was no correlation between the fold changes in gene expression and percentage methylation changes (Figure S1).

### Multivariate analysis of liver high-throughput molecular profiles

We built orthogonal partial least squares discriminant analysis (OPLS-DA) models to assess the predictive ability of the different omics approaches used in this study. The mRNA sequencing and small RNA sequencing, but not the genome methylation data, discriminated the pesticide-treated group from the concurrent control (Table 4).

**Table 3.** Differentially methylated CpG sites located at gene promoters in liver of female Sprague-Dawley rats exposed for 90-days to glyphosate or Roundup MON 52276. Genomic DNA from liver samples were used to perform reduced representation bisulfite sequencing. Differentially methylated CpG sites in gene promoter regions are summarized. Also shown is the consequence on gene expression. (NA, not available; NE, not expressed; NS, not significant). n = 12 per group.

| **DMS for MON 52276** | **qvalue** | **meth.diff** | **gene.name** | **RNA** |
| --- | --- | --- | --- | --- |
| chr10:97895949 (-) | 1.00E-55 | 13.5 | WD repeat domain, phosphoinositide interacting 1 | NS |
| chr2:88365913 (-) | 2.00E-47 | 15.7 | E2F transcription factor 5 | NS |
| chr3:165740932 (-) | 3.00E-43 | 10.4 | zinc finger protein 64 | NS |
| chr1:198181533 (-) | 3.00E-41 | 10.6 | uncharacterized LOC102551806 | NA |
| chr20:6112412 (+) | 1.00E-38 | 10.7 | patatin-like phospholipase domain containing 1 | NE |
| chr8:111686426 (-) | 4.00E-31 | 12.6 | SRP receptor subunit beta | NE |
| chr1:220466269 (-) | 8.00E-29 | 13.8 | transmembrane protein 151A | NS |
| chr3:172382496 (-) | 4.00E-26 | 12.1 | uncharacterized LOC102548488 | NA |
| chr1:142220288 (+) | 2.00E-20 | 15.1 | uncharacterized LOC102553864 | NA |
| chr6:6323934 (-) | 4.00E-18 | 14.5 | uncharacterized LOC103690379 | NA |
| chr19:54317462 (+) | 2.00E-15 | 11 | interferon regulatory factor 8 | NS |
| chr13:78999367 (+) | 5.00E-13 | -11.3 | solute carrier family 9, member C2 (putative) | NE |
| chr13:90115074 (-) | 8.00E-12 | -10.1 | Cd48 molecule | NS |
| chr15:18918298 (+) | 1.00E-08 | 10.5 | POTE ankyrin domain family member H | NS |
| chr5:22028484 (+) | 2.00E-08 | -18.1 | uncharacterized LOC103692299 | NA |
| chr17:80793952 (+) | 2.00E-08 | 10.8 | uncharacterized LOC102550536 | NA |
| chr5:167091490 (+) | 6.00E-07 | 11.3 | microRNA 34a | NA |
| chr8:49709333 (+) | 5.00E-06 | 10.5 | FXYD domain-containing ion transport regulator 2 | NS |
| chr5:169520975 (+) | 3.00E-04 | -10.7 | chromodomain helicase DNA binding protein 5 | NS |
| chr13:69606045 (+) | 5.00E-04 | 12 | uncharacterized LOC108352553 | NA |
| chr4:157099430 (-) | 9.00E-03 | 11.7 | uncharacterized LOC102553636 | NA |
| **DMS for glyphosate** | **qvalue** | **meth.diff** | **gene.name** |  |
| chr3:57717495 (+) | 2.00E-137 | -16.9 | cytochrome b reductase 1 | NS |
| chr3:14455105 (+) | 1.00E-104 | -12.6 | gelsolin | NS |
| chr3:14455057 (-) | 6.00E-104 | -12.5 | gelsolin | NS |
| chr1:219852329 (-) | 5.00E-74 | 17.6 | leucine rich repeat and fibronectin type III domain containing 4 | NE |
| chr13:90115074 (-) | 3.00E-36 | -17.5 | Cd48 molecule | NS |
| chr15:33120272 (+) | 6.00E-34 | -13.7 | RRAD and GEM like GTPase 2 | NE |
| chr12:16912311 (+) | 3.00E-30 | 12.9 | transmembrane protein 184A | NS |
| chr11:72612213 (-) | 1.00E-27 | -10.8 | 3-hydroxybutyrate dehydrogenase 1 | NS |
| chr5:159427478 (+) | 3.00E-27 | -10 | peptidyl arginine deiminase 2 | NE |
| chr14:83218464 (-) | 1.00E-25 | -12 | DEP domain containing 5, GATOR1 subcomplex subunit | NS |
| chr5:22028484 (+) | 4.00E-20 | -26.8 | uncharacterized LOC103692299 | NA |
| chr13:78999367 (+) | 2.00E-15 | -11.1 | solute carrier family 9, member C2 (putative) | NE |
| chr20:3350268 (+) | 1.00E-13 | 13.7 | alpha tubulin acetyltransferase 1 | NE |
| chr1:226291649 (-) | 2.00E-10 | -11 | myelin regulatory factor | NS |
| chr17:80793952 (+) | 9.00E-08 | 10.3 | uncharacterized LOC102550536 | NA |
| chr1:87044466 (+) | 1.00E-06 | 10.9 | galectin 7 | NE |
| chr1:166470212 (+) | 3.00E-05 | -14 | ArfGAP with RhoGAP domain, ankyrin repeat and PH domain 1 | NS |
| chr14:80401981 (+) | 4.00E-05 | -12.1 | carboxypeptidase Z | NS |
| chr13:69606045 (+) | 2.00E-04 | 12.4 | uncharacterized LOC108352553 | NA |
| chr14:80401945 (+) | 3.00E-04 | -11.9 | carboxypeptidase Z | NS |
| chr3:148103483 (-) | 9.00E-03 | 10.5 | defensin beta 25 | NA |

**Table 4.** Predictive ability of high-throughput omics approaches to evaluate the effects of glyphosate and MON 52276. OPLS-DA models were performed for each set of omics data. We show estimates of the total explained variation (R2X), variations between the different groups (R2Y) and the average prediction capability (Q2). We assessed the significance of our classification using permutation tests. New estimates of R2Y and Q2 values were calculated from this 1,000 times permuted dataset (p-values pR2Y and pQ2 for permuted R2Y and Q2, respectively).

| **Comparison** | **R2X** | **R2Y** | **Q2** | **pR2Y** | **pQ2** |
| --- | --- | --- | --- | --- | --- |
| Glyphosate vs control |  |  |  |  |  |
| mRNA-seq | 0.22 | 0.94 | 0.21 | 0.15 | 0.07 |
| Small RNA-seq | 0.39 | 0.99 | 0.52 | 0.13 | 0.002 |
| Genome-wide methylation | 0.13 | 0.66 | 0.03 | 0.06 | 0.89 |
| MON 52276 vs control |  |  |  |  |  |
| mRNA-seq | 0.44 | 0.99 | 0.64 | 0.22 | 0.001 |
| Small RNA-seq | 0.34 | 0.96 | 0.34 | 0.15 | 0.03 |
| Genome-wide methylation | 0.12 | 0.66 | 0.008 | 0.72 | 0.99 |

### Oxidative damage to DNA from increased rates of apurinic/apyrimidinic sites

Since alterations in gene expression reflected triggering of a DNA damage response (Figure 5, Table 2), which could include oxidative stress, we measured DNA damage in the same liver samples. The rate of apurinic/apyrimidinic sites in DNA was increased by the exposure to glyphosate (Figure 9). This confirms that oxidative stress caused by glyphosate exposure resulted in damage to DNA in liver. Oxidative damage to DNA by glyphosate or MON 52276 was not observed in kidney samples (Figure 9).

**Figure 9.**
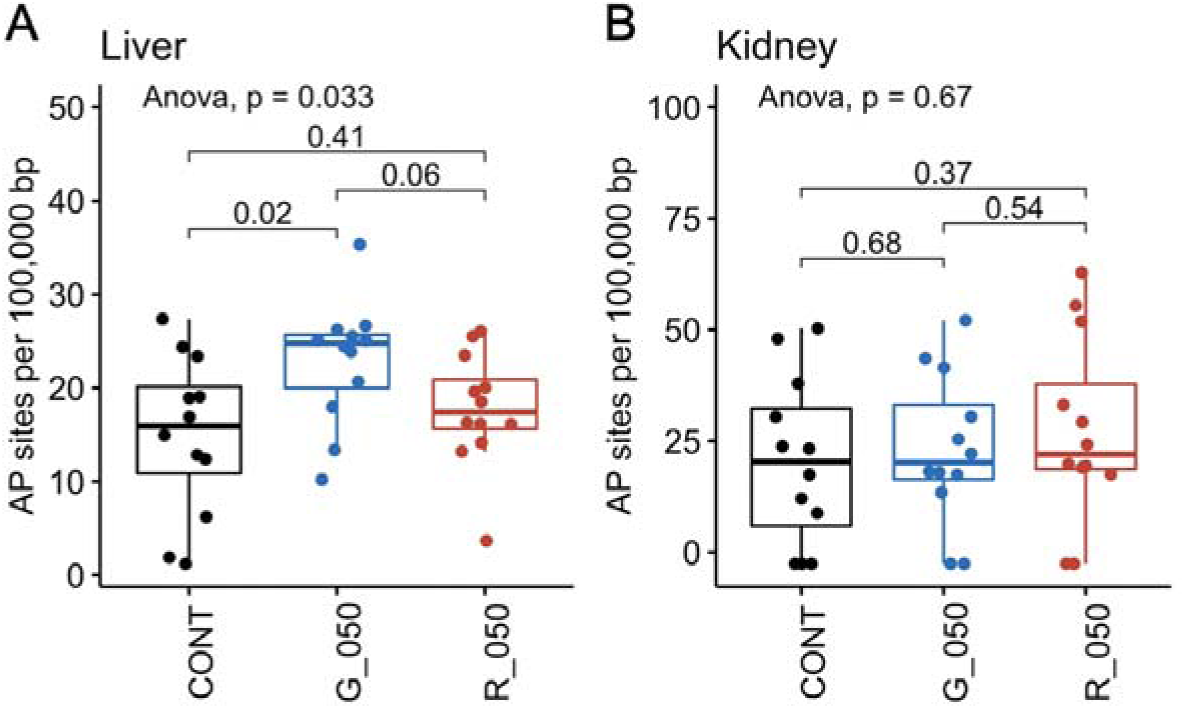
The exposure to glyphosate causes oxidative damage to DNA in rat liver but not in kidneys. DNA from the liver (A) or kidneys (B) of female Sprague-Dawley rats exposed to either 50 mg/kg bw/day glyphosate (G_050) or to an equivalent glyphosate dose of the commercial formulated product Roundup MON 52276 (R_050) for 90-days, was analysed for oxidative damage by measuring the level of apurinic/apyrimidinic (AP) sites compared to untreated animals (CONT). AP site values are shown as box plots with the median, two hinges (the 25th and 75th percentiles), and two whiskers extending to the furthest observation ≤ 1:5 times the interquartile range, along with individual values (solid circles). n = 12 per group. The p-value is from a one-way ANOVA with post-hoc Tukey HSD.

Overall, the *in vivo* toxicogenomic evaluation is in accord with the results obtained from the ToxTracker assay system, which indicate that MON 52276 was causing more biological changes than glyphosate.

## Discussion

Many controversies exist around genotoxic effects of glyphosate and its commercial formulated herbicide products. In an effort to address some of the outstanding issues we compared the biological changes caused by glyphosate and three GBH Roundup formulations, with a focus on the EU representative formulation Roundup MON 52276.

The results presented in this study confirm the increased incidence of lesions (steatosis, necrosis) in the liver of animals treated with MON 52276 (Figure 3; Table 1), which was suggested from the analysis of the serum metabolite profile described in our previous publication (Mesnage et al. 2021). The finding of liver steatosis from exposure to MON 52276 is also in agreement with our observation that an ultra-low dose of Roundup Grand Travaux Plus administered to the same strain of Sprague-Dawley rats over a 2-year period resulted in non-alcoholic fatty liver disease (Mesnage et al. 2015, Mesnage et al. 2017). However, the exposure time was not suitable to detect the development of chronic disease and long-term, ideally life-long studies will be necessary to understand whether glyphosate can cause organ damage, which may potentially be associated with cancer development. Although in this study glyphosate alone only caused limited changes in liver molecular and histological profiles, longer studies would be needed to understand if the non-statistically significant increases in signs of organ damage (Table 1) could lead to adverse health effects if exposure was prolonged.

Alterations in liver gene expression and small RNA, particularly miRNA profiles observed in this study provide insights into the mode-of-action of glyphosate as a carcinogen. Glyphosate and Roundup MON 52276 gave rise to a consistent alteration in the levels of expression of 9 genes in liver (Table 2) with some also similarly altered in kidney (Figure 6) whose function is linked with an oxidative stress and DNA damage response with possible carcinogenic consequences. In addition, as the same alterations in gene expression were seen with both glyphosate and MON 52276, our findings suggest these changes in gene function are largely due to glyphosate and not the co-formulants present in the commercial herbicide formulation. However, a limitation of this study is that connections between the omics results and physiological/pathological presentations are insufficient to fully confirm that the molecular changes observed can be used as biomarkers. There was no evidence of liver tumors in female rats in this study, although some studies in rodents have shown development of liver adenomas in male Sprague-Dawley rats and hepatocellular adenomas in male Wistar rats following chronic exposure (Portier 2020). A new study evaluating whether changes in molecular profiles in a young animal can predict organ damage in late life would be appropriate to confirm the relevance of these potential markers of toxicity. Furthermore, the fact that non-alcoholic fatty liver disease (NAFLD) is an established major risk factor for liver cancer (Massoud and Charlton 2018) is noteworthy, since our observation in this (Table 1; Figure 3) and our previous study (Mesnage et al. 2017) show that Roundup exposure can result in NAFLD. This suggests that GBH exposure may in some cases contribute to the development of NAFLD and thus predispose individuals to developing liver cancer. Such an indirect role of GBH exposure to liver cancer predisposition would account why to date this is not a recognised contributory risk factor in this disease.

We show that glyphosate caused oxidative DNA damage in liver by increasing the rate of formation of apurinic/apyrimidinic DNA sites (Figure 9). This could be linked with the alteration in the expression of genes associated with the induction/repair of DNA damage via an alteration in TP53 signaling, which we observe in our transcriptome analysis (Figure 4; Table 2). However, no genotoxic activity was detected in the 6 ToxTracker mES reporter cell lines for glyphosate (Figure 2), which indicates that glyphosate does not act as a direct genotoxicant or a mutagen. These data taken together suggest that DNA damage from glyphosate or MON 52276 exposure could be the result of organ damage from oxidative stress and concomitant inflammatory processes, which can be induced at least in part by the observed fatty liver condition as well as necrosis.

The miRNAs, which had their levels altered in liver samples by glyphosate and Roundup MON 52276 (Figure 7B), have been shown to be involved in carcinogenesis. The level of miR-22, which was decreased by MON 52276 treatment, has tumor suppressor properties by modulating p53-dependent cellular fate (Tsuchiya et al. 2011). The transactivation of miR-22 occurs in response to DNA damage to repress p21 expression and activate apoptosis (Sun et al. 2019). The levels of miR-17 were also decreased by MON 52276 and miR-17 is known to interact with a p53 activating kinase and forms a positive feed-forward loop to promote tumorigenesis (Cai et al. 2015). Although glyphosate did not alter the levels of miR-22 and miR-17, it did give rise to a decrease in miR-30 and an increase in miR-10 (Figure 7B). Both miR-30 and miR-10 also have known links with cancer processes. MiR-30 regulates mitochondrial fission by targeting p53 (Li et al. 2010) and has been proposed to be an “oncomiR” in gastric cancer cells (Wang, Jiao et al. 2017) whereas miR-10 has been found to be increased in several types of leukaemia (Garzon et al. 2008). Collectively, the alterations in the levels of these miRNAs support the involvement of the p53 pathway in DNA damage caused by glyphosate or MON 52276 with possible carcinogenic outcomes.

An increasing number of studies have found that circulating small RNAs can serve as biomarkers for early diagnosis of lymphomas. Serum miR-22 is a non-invasive predictor of poor clinical outcome in patients with diffuse large B-cell lymphoma (Marchesi et al. 2018). This suggests that future studies measuring these miRNAs in blood could act as a biomarker for environmental epidemiology studies measuring effects from glyphosate and GBH exposure (Jorgensen et al. 2020). In addition, the genes which had their expression consistently altered by both glyphosate and Roundup MON 52276 (Table 2) can serve as a biomarker of glyphosate effects on p53 cellular pathways. Molecular changes of this type detected at early stages of carcinogenesis while no pathological effects are observable are increasingly used to predict long-term carcinogenesis (Locke et al. 2019).

Other studies have suggested that glyphosate causes oxidative damage to DNA, and that this can activate DNA repair mechanisms. A study on human peripheral blood mononuclear cells showed that glyphosate decreased the expression of the tumor suppressor gene *TP53,* and that this decreased expression was concomitant with an induction of 5-mC methylation of the *TP53* promoter (Woźniak et al. 2020). In addition, changes in the expression of the core-clock gene *NR1D1,* whose expression was most affected by the exposure to glyphosate in our study, are known to impact the circadian phenotype of *TP53*, ultimately affecting proliferation, apoptosis, and cell migration (Basti et al. 2020). Interestingly, disturbances of circadian genes are known to be involved in the etiology of Non-Hodgkin lymphoma (NHL) (Hoffman et al. 2009), providing a molecular mechanistic explanation for observations stemming from epidemiological studies showing that night-time work predisposes to NHL (Lahti et al. 2008). The circadian clock is known to impact cell proliferation and migration in lymphoma cells (Abreu et al. 2018). However, known epigenomic differences between tissues imply that these findings may be specific to liver and kidney and that other organs and cell systems would have to be studied to assess if the molecular biological changes we observed from glyphosate and MON 52276 exposure have similar effects in other tissues.

Oxidative damage to DNA reflected by the level of apurinic/apyrimidinic (AP) sites was observed with glyphosate but not with MON 52276, and only in liver (Figure 9). One possible explanation for this apparent contradiction is that the damage caused by MON 52276 was sufficiently high to elicit DNA repair, while DNA damage for glyphosate was not detected by DNA lesion sensor mechanisms. This is corroborated by the observation of TP53 pathway activation with MON 52276 but not with glyphosate. It is thus possible that MON 52276 caused oxidative DNA damage, but this could not be detected because the degree of damage is low enough to be repaired.

The findings from our comparative toxicogenomic evaluation in rats suggested that MON 52276 caused oxidative DNA damage. None of the GFP-reporters from the ToxTracker assay system scored positive for activation of a DNA damage response or p53-mediated cellular stress (Figure 2). However, the Srxn1-GFP reporter for oxidative stress was activated upon exposure to MON 52276 suggesting that oxidative stress observed after 24h of exposure could be driving DNA damage long term, in line with the in vivo data. This also suggests that there are some other physiological factors which can influence glyphosate genotoxicity. This is not necessarily surprising given the large number of studies with contradictory results and divergent interpretations (Benbrook 2019). Even if the surfactants in the formulations can exert cytotoxic effects by altering cell membrane structure, it is also possible that glyphosate uptake *in vitro* is limited, as glyphosate is hydrophilic and enters cells via L-type amino acid transporters (Xu et al. 2016). The observation in an *in vitro* study, which showed that ∼ 20% of ^14^C-glyphosate entered HepG2 cells is in support of this possibility (Gasnier et al. 2011). It is also possible that DNA damage is the consequence of the activation of another type of toxicity process, which would not be detectable by the ToxTracker assay. For instance, DNA damage could be secondary to liver inflammation or oxidative stress (Kawanishi et al. 2017).

MON 52276 was found to be the most potent at the gene expression and phenotypic-level, though glyphosate caused greatest changes in CpG methylation. However, it is not clear to which extent the analysis of the number of differentially methylated CpG sites is robust. Our analysis of the predictive ability of this genome-wide methylation data showed that it was not able to discriminate the pesticide-treated groups from the concurrent control by contrast to the gene expression data (Table 4). This suggests that our experimental design is underpowered to detect changes in genome-wide methylation, which probably originate in the intrinsic properties of DNA methylation data. By comparison to gene expression analysis, which uses quantitative count data, CpG methylation involves the analysis of a larger number of variables, which are semi-quantitative as they are essentially a ratio between the number of C bases that are methylated and the number of C bases that are unmethylated. This is thus more subject to experimental variation. Even if the biological significance of the observed changes in liver DNA methylation profiles (Figure 7) remain unclear, our findings constitute a first step towards the creation of a biomarker allowing the identification of glyphosate-associated pathologies in human populations. Although other epigenetics studies have been performed in different model systems (Duforestel et al. 2019, Kubsad et al. 2019, Woźniak et al. 2020), no *in vivo* biomarker of glyphosate exposure has been proposed or identified in laboratory animals. The increase in Tet3 activity, which was proposed to be a target of glyphosate in the mammary gland (Duforestel et al. 2019), was unchanged in our study. This could be due to tissue specific effects, or to the regulation of Tet3 activity at a protein rather than gene transcription level, which would not be detectable in our study. Since cancer risks are associated with genomic instability, early events in tumorigenesis can be reflected by epigenetic changes (Locke et al. 2019). For instance, the study of DNA methylation levels in Long Interspersed Nucleotide Element 1 (LINE-1) was found to be a surrogate for changes in global DNA methylation levels caused by exposure to pesticides in farmers (Alexander et al. 2017). Epigenetic biomarkers can also be more specific as in the case of the methylation levels of *WRAP53*, which is antisense to *TP53,* was found to be associated to pesticide exposure in a Mexican population (Paredes-Céspedes et al. 2019). Epigenetic signatures can also be compound-specific such as for smoking, and even inform on smoking habits and time since smoking cessation (Kaur et al. 2019).

While the co-formulants contained in the representative EU commercial GBH formulation MON 52276 are known to have one of the best safety profiles among agricultural surfactants, the results from the study presented here and in our previous investigation with the same animals (Mesnage et al. 2021) suggests that it is more toxic than previously claimed. Previous studies on human cells found that MON 52276 was no more cytotoxic than glyphosate alone at equivalent concentrations (Mesnage et al. 2013), and that it was less toxic than glyphosate on rainbow trout and water fleas (Mesnage et al. 2019). However, our study shows that exposure to MON 52276 causes liver steatosis and necrosis and supports the need for long-term toxicity evaluations of the toxicity of formulated products. MON 76207 could be even more cytotoxic as suggested by cytotoxicity profiles in mES cells (Figure S2). Since the residues of co-formulants are not monitored in human populations or in the environment, and the fact that there is no regulatory requirement to assess their toxicity in long-term studies, their health risks remain a major unknown quantity (Mesnage et al. 2019). Only a few studies have shown that the most used co-formulant POEA can persist in the environment after agricultural applications (Tush et al. 2018). No studies have been performed to understand if pre-harvest applications of GBH, particularly of cereal food crops, could expose consumers to co-formulant residues. Professional pesticide applicators are always exposed to formulated GBH products or mixed with additional pesticides, even if biomonitoring data is needed to understand internal exposures resulting from the application of agricultural sprays.

Farmers frequently mix pesticides in order to overcome weed resistance to a single herbicide. Glyphosate is increasingly applied in combination with other active ingredients such as dicamba, mesotrione, metolachlor or 2,4-D in order to mitigate herbicide-resistance in large-scale monocultures of herbicide-tolerant GM crops (Vink et al. 2012, Bonny 2016). An increasing number of studies have found that pesticides can exert effects as mixtures at concentrations, which they do not show negative health outcomes in isolation (Martin et al. 2021). Future studies will have to evaluate the combined effects of glyphosate with other pesticides, since human populations are rarely exposed to single compounds.

## Conclusions

Our study reports the first observations of the changes in the epigenome (DNA methylation profile), transcriptome and miRNA profiles in the liver of rats exposed to glyphosate or its formulated product Roundup MON 52276. We have confirmed that glyphosate causes DNA damage and leads to activation of DNA repair mechanisms in a mammalian model system *in vivo*. Mechanistic evaluation of carcinogenicity mode-of-action showed that Roundup herbicides can activate oxidative stress and an unfolded protein response at concentrations at which glyphosate gave no effects. Our study further highlights the power of high-throughput ‘omics’ methods to detect metabolic changes, which are poorly reflected or completely missed within conventional biochemical and histopathological measurements upon which regulators currently rely. Thus, adoption by regulatory agencies of multi-omics analyses would result in more accurate evaluation of a chemical’s toxicity and therefore better protection measures being enacted with major public health benefits.

## Acknowledgements

This work was funded by the Sustainable Food Alliance (USA), the Heartland Health Research Alliance (USA) and in part by the Sheepdrove Trust (UK), all of whose support is gratefully acknowledged.

---

**Figure S1.**
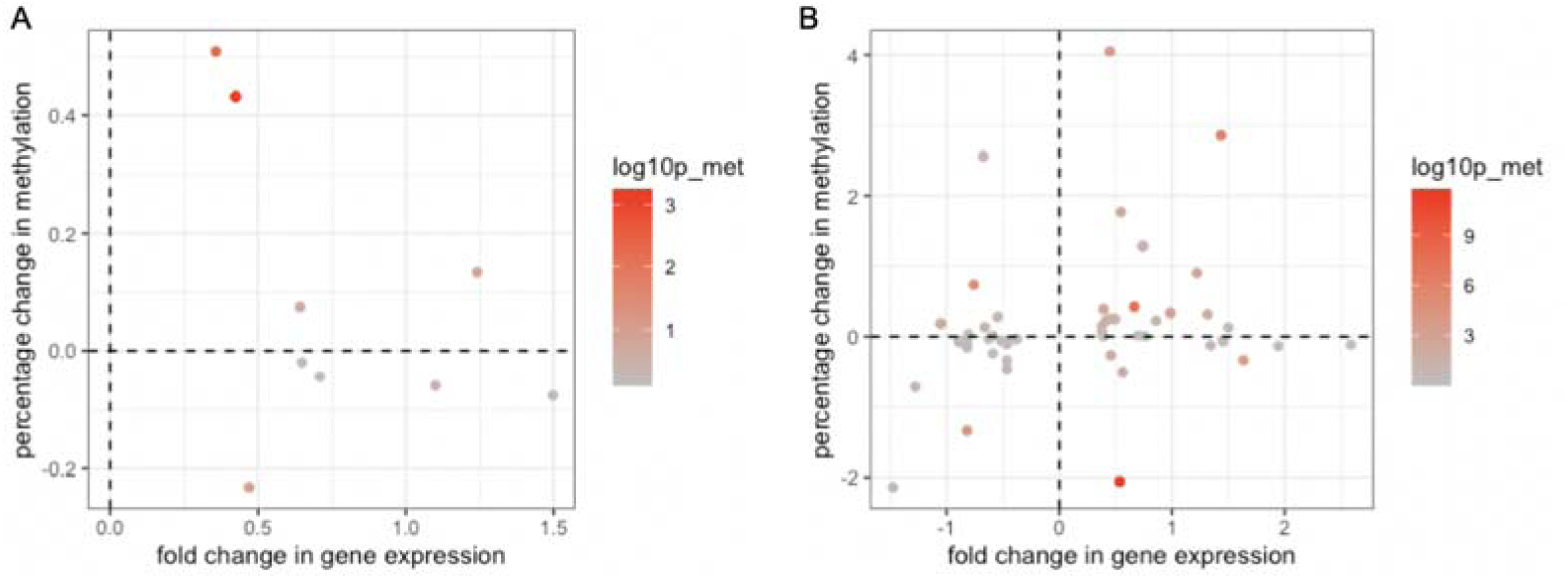
No correlation between the fold changes in gene expression and percentage methylation changes. RRBS of DNA from female Sprague-Dawley rat liver samples was performed to assess if alterations in epigenetic (DNA methylation) status may be responsible for the glyphosate (**A**) and MON 52276 (**B**)-related changes in gene expression patterns. The fold changes in gene expression (RNA-seq) are compared to the percentage of change in methylation (RRBS) for the differentially expressed genes.

**Figure S2.**
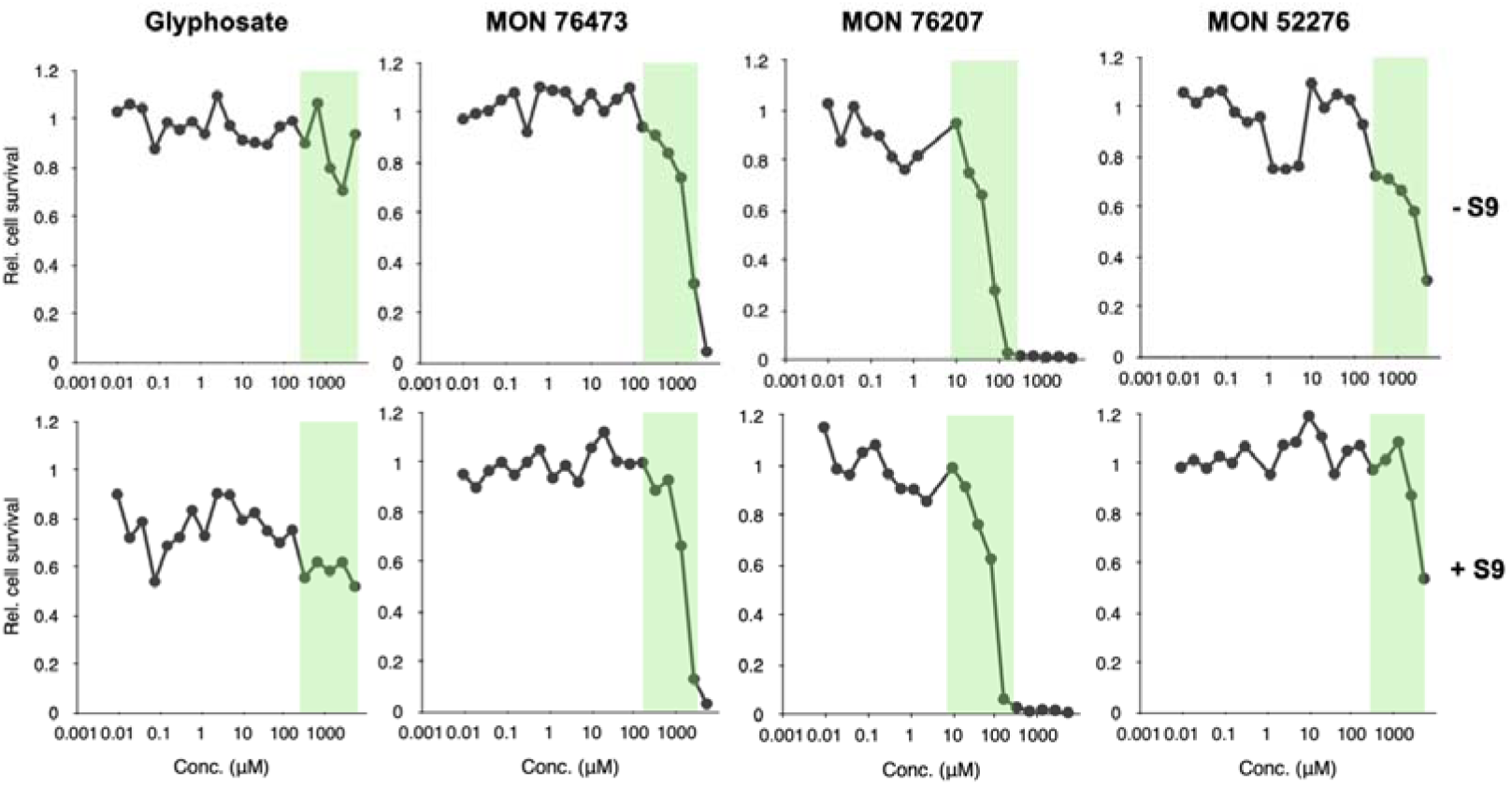
Cytotoxicity dose response evaluation of glyphosate and 3 Roundup herbicides in mouse embryonic stem cells. A total of 20 concentrations were tested for each compound to determine cytotoxicity thresholds. The green shaded area denotes the selected concentrations used in the ToxTracker assay system.

**Figure S3.**
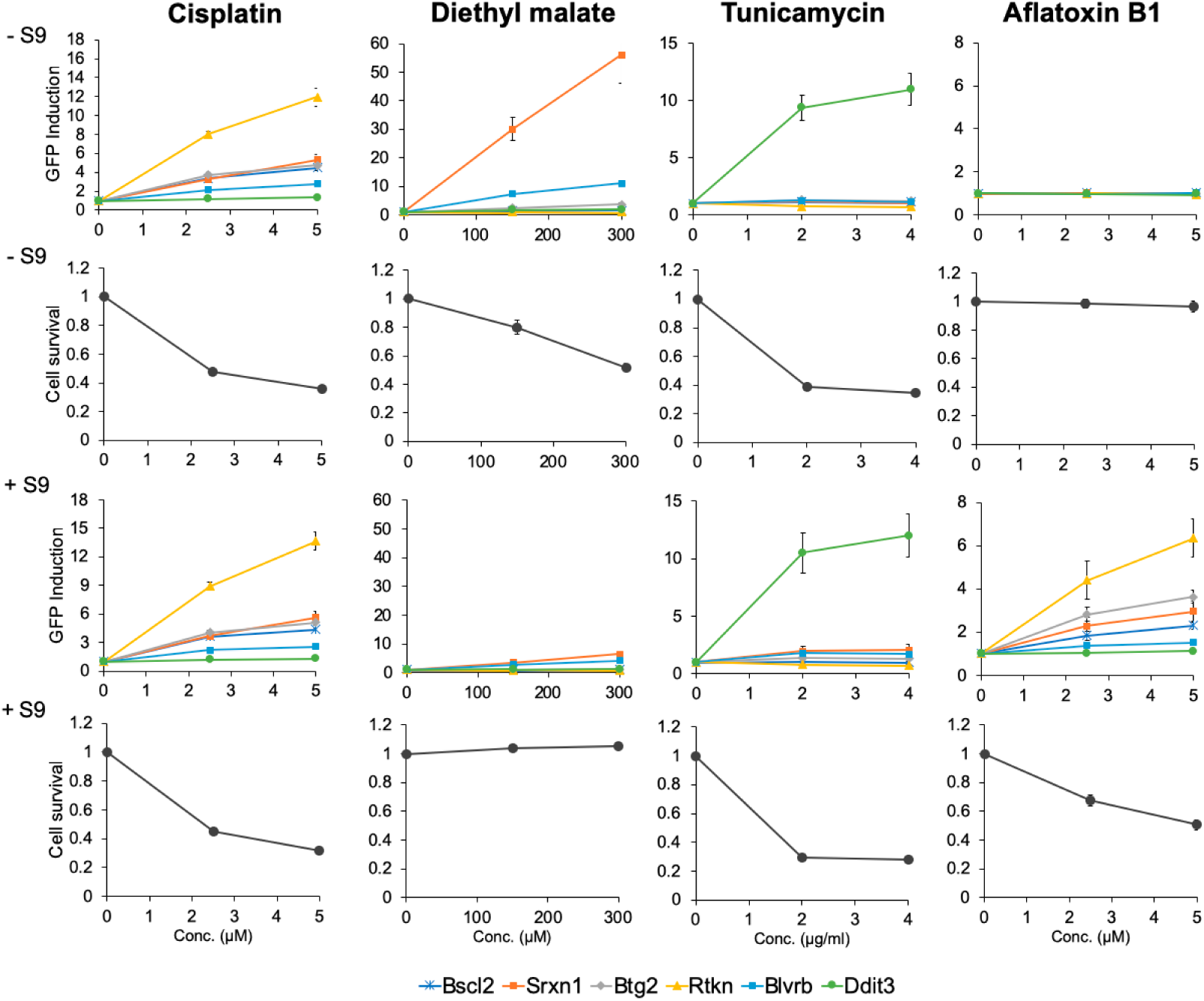
Positive reference treatments with cisplatin (DNA damage), diethyl maleate (oxidative stress), tunicamycin (unfolded/misfolded protein response) and aflatoxin B1 (metabolic activation of progenotoxins by S9) were included in all the ToxTracker system experiments.

**Table S1.** Food consumption in adult female Sprague-Dawley rats administered for 90-days with glyphosate and Roundup MON 52276. No differences were found. n = 12 per group.

| Consumption / Groups | Control | Glyphosate |  |  | MON 52276 |  |  |
| --- | --- | --- | --- | --- | --- | --- | --- |
| Dose (mg/kg) | 0 | 0.5 | 50 | 175 | 0.5 | 50 | 175 |
| Week 0 | 16.3 ± 2.1 | 18.3 ± 3.6 | 19.6 ± 1.6 | 21.3 ± 1.6 | 18.3 ± 3.3 | 16.7 ± 4.9 | 17.5 ± 3.5 |
| Week 1 | 22.5 ± 2.2 | 20.8 ± 2.2 | 24.2 ± 1 | 23.3 ± 1.4 | 24.2 ± 2.9 | 22.5 ± 2.9 | 22.5 ± 2.2 |
| Week 2 | 22.1 ± 1.6 | 21.3 ± 0.8 | 21.3 ± 1.6 | 20.8 ± 2.2 | 21.3 ± 0.8 | 20.4 ± 0.8 | 20 ± 1.4 |
| Week 3 | 20.8 ± 2.2 | 19.2 ± 1 | 18.3 ± 1.9 | 17.9 ± 2.5 | 16.7 ± 1.9 | 16.3 ± 2.5 | 17.5 ± 1 |
| Week 4 | 18.3 ± 1.9 | 17.9 ± 0.8 | 17.9 ± 0.8 | 17.5 ± 1 | 19.6 ± 0.8 | 17.1 ± 1.6 | 17.1 ± 1.6 |
| Week 5 | 19.2 ± 2.9 | 17.5 ± 3.5 | 18.3 ± 1.4 | 19.6 ± 1.6 | 20 ± 2.4 | 18.8 ± 1.6 | 17.1 ± 1.6 |
| Week 6 | 20.4 ± 1.6 | 20.4 ± 0.8 | 20.4 ± 0.8 | 20 ± 1.4 | 20.4 ± 0.8 | 17.9 ± 2.5 | 17.5 ± 1 |
| Week 7 | 24.2 ± 1 | 21.3 ± 1.6 | 20.4 ± 2.8 | 18.8 ± 2.5 | 20 ± 1.4 | 18.8 ± 2.8 | 18.3 ± 1.4 |
| Week 8 | 18.3 ± 2.4 | 19.2 ± 2.9 | 19.2 ± 1.7 | 19.6 ± 0.8 | 18.8 ± 1.6 | 18.8 ± 0.8 | 17.5 ± 1 |
| Week 9 | 22.1 ± 2.8 | 20.8 ± 2.2 | 22.1 ± 1.6 | 22.5 ± 1 | 22.9 ± 1.6 | 22.9 ± 2.5 | 22.5 ± 1.7 |
| Week 10 | 20.4 ± 1.6 | 17.5 ± 1 | 19.2 ± 2.2 | 20.4 ± 0.8 | 19.2 ± 1.7 | 19.6 ± 0.8 | 17.9 ± 1.6 |
| Week 11 | 19.2 ± 2.9 | 18.3 ± 1.4 | 20 ± 1.4 | 18.8 ± 1.6 | 19.2 ± 1 | 17.9 ± 1.6 | 18.8 ± 1.6 |
| Week 12 | 19.6 ± 1.6 | 20 ± 1.4 | 20 ± 1.4 | 18.8 ± 0.8 | 18.8 ± 1.6 | 17.5 ± 1 | 18.3 ± 1.4 |
| Week 13 | 20 ± 2.4 | 20.4 ± 2.1 | 22.1 ± 1.6 | 20.4 ± 0.8 | 21.3 ± 2.1 | 20.4 ± 0.8 | 20 ± 2.4 |
| Mean | 20.5 ± 1.8 | 19.6 ± 0.8 | 20.3 ± 0.5 | 19.6 ± 0.6 | 20.2 ± 0.8 | 19.1 ± 1.3 | 18.8 ± 0.8 |

**Table S2.**
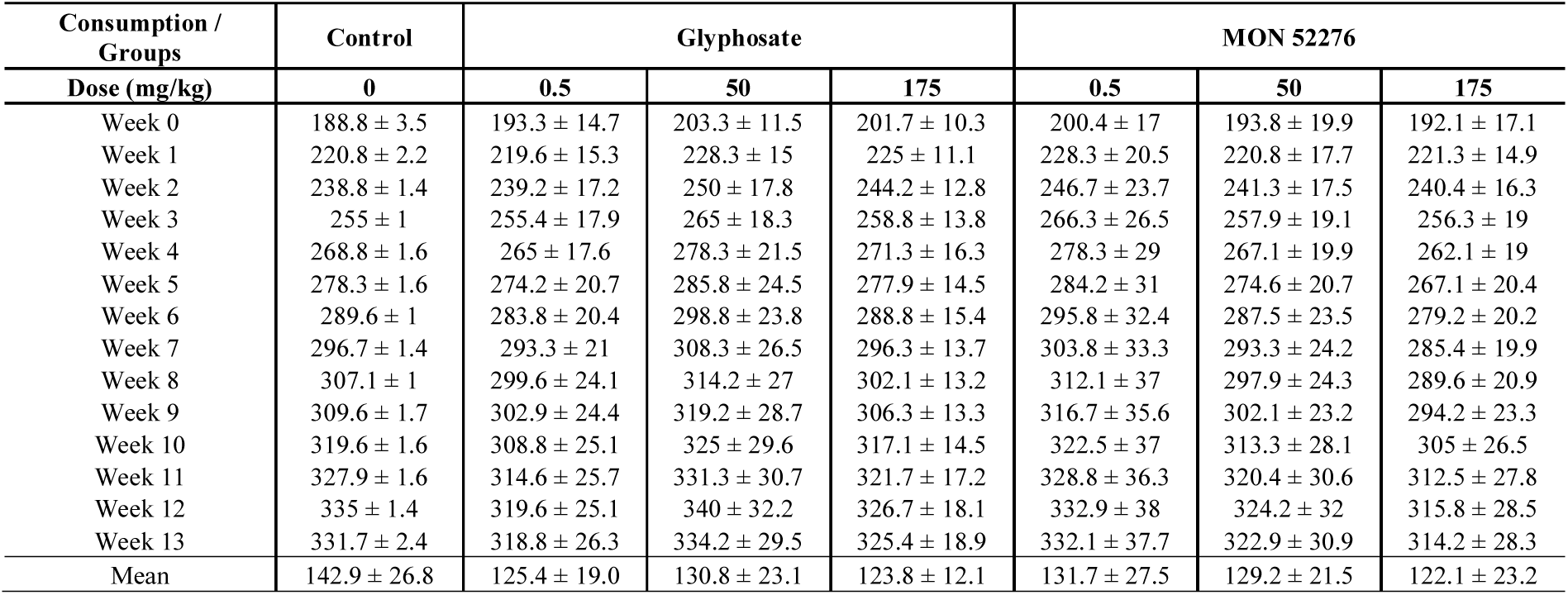
Body weights in adult female Sprague-Dawley rats administered for 90-days with glyphosate and Roundup MON 52276. No differences were found. n = 12 per group.

| Consumption / Groups | Control | Glyphosate |  |  | MON 52276 |  |  |
| --- | --- | --- | --- | --- | --- | --- | --- |
| Dose (mg/kg) | 0 | 0.5 | 50 | 175 | 0.5 | 50 | 175 |
| Week 0 | 188.8 ± 3.5 | 193.3 ± 14.7 | 203.3 ± 11.5 | 201.7 ± 10.3 | 200.4 ± 17 | 193.8 ± 19.9 | 192.1 ± 17.1 |
| Week 1 | 220.8 ± 2.2 | 219.6 ± 15.3 | 228.3 ± 15 | 225 ± 11.1 | 228.3 ± 20.5 | 220.8 ± 17.7 | 221.3 ± 14.9 |
| Week 2 | 238.8 ± 1.4 | 239.2 ± 17.2 | 250 ± 17.8 | 244.2 ± 12.8 | 246.7 ± 23.7 | 241.3 ± 17.5 | 240.4 ± 16.3 |
| Week 3 | 255 ± 1 | 255.4 ± 17.9 | 265 ± 18.3 | 258.8 ± 13.8 | 266.3 ± 26.5 | 257.9 ± 19.1 | 256.3 ± 19 |
| Week 4 | 268.8 ± 1.6 | 265 ± 17.6 | 278.3 ± 21.5 | 271.3 ± 16.3 | 278.3 ± 29 | 267.1 ± 19.9 | 262.1 ± 19 |
| Week 5 | 278.3 ± 1.6 | 274.2 ± 20.7 | 285.8 ± 24.5 | 277.9 ± 14.5 | 284.2 ± 31 | 274.6 ± 20.7 | 267.1 ± 20.4 |
| Week 6 | 289.6 ± 1 | 283.8 ± 20.4 | 298.8 ± 23.8 | 288.8 ± 15.4 | 295.8 ± 32.4 | 287.5 ± 23.5 | 279.2 ± 20.2 |
| Week 7 | 296.7 ± 1.4 | 293.3 ± 21 | 308.3 ± 26.5 | 296.3 ± 13.7 | 303.8 ± 33.3 | 293.3 ± 24.2 | 285.4 ± 19.9 |
| Week 8 | 307.1 ± 1 | 299.6 ± 24.1 | 314.2 ± 27 | 302.1 ± 13.2 | 312.1 ± 37 | 297.9 ± 24.3 | 289.6 ± 20.9 |
| Week 9 | 309.6 ± 1.7 | 302.9 ± 24.4 | 319.2 ± 28.7 | 306.3 ± 13.3 | 316.7 ± 35.6 | 302.1 ± 23.2 | 294.2 ± 23.3 |
| Week 10 | 319.6 ± 1.6 | 308.8 ± 25.1 | 325 ± 29.6 | 317.1 ± 14.5 | 322.5 ± 37 | 313.3 ± 28.1 | 305 ± 26.5 |
| Week 11 | 327.9 ± 1.6 | 314.6 ± 25.7 | 331.3 ± 30.7 | 321.7 ± 17.2 | 328.8 ± 36.3 | 320.4 ± 30.6 | 312.5 ± 27.8 |
| Week 12 | 335 ± 1.4 | 319.6 ± 25.1 | 340 ± 32.2 | 326.7 ± 18.1 | 332.9 ± 38 | 324.2 ± 32 | 315.8 ± 28.5 |
| Week 13 | 331.7 ± 2.4 | 318.8 ± 26.3 | 334.2 ± 29.5 | 325.4 ± 18.9 | 332.1 ± 37.7 | 322.9 ± 30.9 | 314.2 ± 28.3 |
| Mean | 142.9 ± 26.8 | 125.4 ± 19.0 | 130.8 ± 23.1 | 123.8 ± 12.1 | 131.7 ± 27.5 | 129.2 ± 21.5 | 122.1 ± 23.2 |

**Table S3.** Water consumption in adult female Sprague-Dawley rats administered for 90-days with glyphosate and Roundup MON 52276. Statistical significance is shown for Dunnett’s test on mean consumption (**, P≤0.01). n = 12 per group.

| Consumption / Groups | Control | Glyphosate |  |  | MON 52276 |  |  |
| --- | --- | --- | --- | --- | --- | --- | --- |
| Dose (mg/kg) | 0 | 0.5 | 50 | 175 | 0.5 | 50 | 175 |
| Week 0 | 22.5 ± 2.5 | 24.2 ± 4.4 | 26.5 ± 2.8 | 26.8 ± 3.9 | 26.7 ± 5.9 | 24.5 ± 5.4 | 25.5 ± 6 |
| Week 1 | 30.5 ± 3.6 | 30.3 ± 1.6 | 29.7 ± 0.9 | 28.3 ± 2.2 | 32.5 ± 5.5 | 28.5 ± 1 | 24.3 ± 3.9 |
| Week 2 | 29.2 ± 5.1 | 28.2 ± 2.4 | 26 ± 0.5 | 27.3 ± 1.5 | 28.8 ± 3 | 26.3 ± 3.1 | 18.7 ± 4.2 |
| Week 3 | 27.5 ± 0.8 | 25.5 ± 4.8 | 24.3 ± 1.8 | 22.8 ± 2.7 | 26.8 ± 4.9 | 23.5 ± 1.5 | 17 ± 2.6 |
| Week 4 | 22.3 ± 2 | 23.1 ± 0.7 | 21.7 ± 1.8 | 20 ± 1.7 | 22.5 ± 2.9 | 22 ± 1.8 | 15 ± 1.4 |
| Week 5 | 25.2 ± 1.5 | 24.7 ± 3.6 | 25.2 ± 1.5 | 23 ± 2.7 | 26.3 ± 2.7 | 25.3 ± 2.6 | 13.5 ± 2.3 |
| Week 6 | 25.7 ± 1.6 | 26.5 ± 3.9 | 23.7 ± 1.4 | 23.5 ± 3 | 27.2 ± 3.9 | 23.5 ± 2.3 | 17.5 ± 1.4 |
| Week 7 | 25.5 ± 2.5 | 24.7 ± 2.4 | 23 ± 1.9 | 21 ± 2.3 | 24.8 ± 4.1 | 20.8 ± 1.4 | 17.3 ± 2.1 |
| Week 8 | 24.5 ± 3.3 | 25 ± 3.4 | 26 ± 2.8 | 22.2 ± 3.8 | 24 ± 4 | 23.2 ± 0.6 | 18.8 ± 1.8 |
| Week 9 | 26.3 ± 4.3 | 24.3 ± 2.5 | 25.3 ± 1.2 | 23.2 ± 2.2 | 25.7 ± 2.1 | 25.2 ± 2.3 | 17.2 ± 4.4 |
| Week 10 | 27.7 ± 1.6 | 24.5 ± 1.9 | 24.3 ± 2.9 | 22 ± 3.6 | 27 ± 4 | 23.8 ± 0.8 | 17.5 ± 1.8 |
| Week 11 | 25.2 ± 2.6 | 24.5 ± 1.8 | 24.7 ± 1.6 | 19.7 ± 3.2 | 25.5 ± 3 | 23.5 ± 1.5 | 16.5 ± 0.8 |
| Week 12 | 28.5 ± 2.1 | 30.2 ± 2.7 | 26.8 ± 1.1 | 20 ± 2 | 27.3 ± 3.5 | 23.8 ± 2.8 | 17.5 ± 1.3 |
| Week 13 | 27.5 ± 2.9 | 29.2 ± 3.5 | 30.2 ± 2.1 | 25.7 ± 1.8 | 31.0 ± 3.0 | 29.0 ± 2.0 | 19.8 ± 2.5 |
| Mean | 26.6 ± 2.2 | 26.2 ± 1.9 | 25.4 ± 1.3 | 23.0 ± 1.8 | 26.9 ± 3.2 | 24.5 ± 0.4 | 17.7 ± 1.8 ** |

**Table S4.**
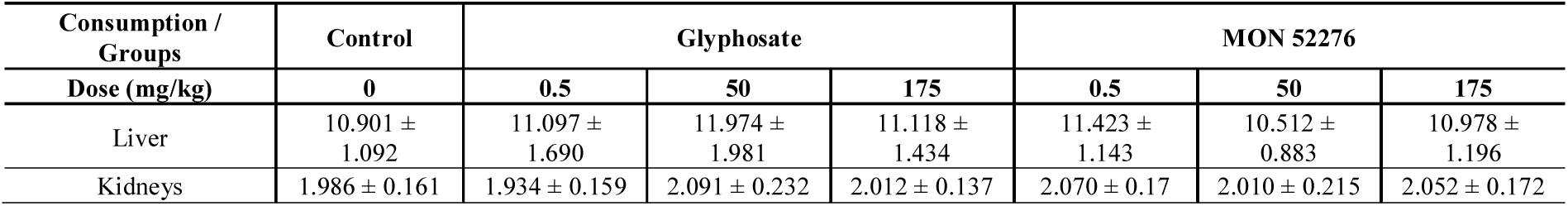
Liver and kidney weights in adult female Sprague-Dawley rats administered for 90-days with glyphosate and Roundup MON 52276. No differences were found. n = 12 per group.

| Consumption / Groups | Control | Glyphosate |  |  | MON 52276 |  |  |
| --- | --- | --- | --- | --- | --- | --- | --- |
| Dose (mg/kg) | 0 | 0.5 | 50 | 175 | 0.5 | 50 | 175 |
| Liver | 10.901 ± 1.092 | 11.097 ± 1.690 | 11.974 ± 1.981 | 11.118 ± 1.434 | 11.423 ± 1.143 | 10.512 ± 0.883 | 10.978 ± 1.196 |
| Kidneys | 1.986 ± 0.161 | 1.934 ± 0.159 | 2.091 ± 0.232 | 2.012 ± 0.137 | 2.070 ± 0.17 | 2.010 ± 0.215 | 2.052 ± 0.172 |

